# Tuning Extracellular Electron Transfer by *Shewanella oneidensis* Using Transcriptional Logic Gates

**DOI:** 10.1101/2019.12.18.881623

**Authors:** Christopher M. Dundas, Benjamin K. Keitz

## Abstract

Extracellular electron transfer pathways, such as those in the bacterium *Shewanella oneidensis*, interface cellular metabolism with a variety of redox-driven applications. However, designer control over EET flux in *S. oneidensis* has proven challenging since a functional understanding of its EET pathway proteins and their effect on engineering parameterizations (e.g., response curves, dynamic range) is generally lacking. To address this, we systematically altered transcription and translation of single genes encoding parts of the primary EET pathway of *S. oneidensis*, CymA/MtrCAB, and examined how expression differences affected model-fitted parameters for Fe(III) reduction kinetics. Using a suite of plasmid-based inducible circuits maintained by appropriate *S. oneidensis* knockout strains, we pinpointed construct/strain pairings that expressed *cymA, mtrA*, and *mtrC* with maximal dynamic range of Fe(III) reduction rate. These optimized EET gene constructs were employed to create Buffer and NOT gate architectures, that predictably turn on and turn off EET flux, respectively, in response to IPTG. Furthermore, we found that response functions generated by these logic gates (i.e., EET activity vs. inducer concentration) were comparable to those generated by conventional synthetic biology circuits, where fluorescent reporters are the output. Our results provide insight on programming EET activity with transcriptional logic gates and suggest that previously developed transcriptional circuitry can be adapted to predictably control EET flux.

## Introduction

Extracellular electron transfer (EET) is an anaerobic respiratory mechanism through which electroactive microbes interact with and influence their environment.^1^ For example, EET flux can be harnessed for power generation in microbial fuel cells,^2^ the remediation of environmental pollutants,^3^ and for material synthesis.^4,5^ In theory, EET can connect biological computation with modern electronics, potentially enabling the creation of new sensors and other hybrid living-electronic devices.^6^ Because redox-active proteins and metabolites mediate microbial electron transfer, these applications have the potential for genetic and metabolic optimization. However, strict control over expression/activity of metabolic and proteomic elements governing EET flux and thus programmable control over cellular electron flux remains challenging.

*Shewanella oneidnesis* is a facultative bacterium that couples carbon oxidation to EET flux^7^. The EET pathways of *S. oneidensis* are common engineering targets, owing to the bacteria’s favorable growth characteristics and genetic tractability. In the absence of oxygen, these pathways facilitate respiration onto a variety of technologically-relevant metal species, including Fe(III),^8^ Mn(IV), and Pd(II).^9^ The primary EET pathway in *S. oneidensis* is the CymA/MtrCAB conduit (Figure 1), which is comprised of a periplasmic/outer membrane *c-*cytochrome network that links carbon metabolism to the reduction of inorganic substrates.^10^ Manipulating EET in *S. oneidensis* has been predominantly explored in the context of bioelectrochemical applications.^11^ For example, a frequent power generation goal to increase EET flux above wild-type *S. oneidensis* levels can be accomplished through overexpression of flavin biosynthesis enzymes.^12^ However, controlled expression of the CymA/Mtr components has received relatively little attention outside of electrode-based characterizations.

**Figure 1.**
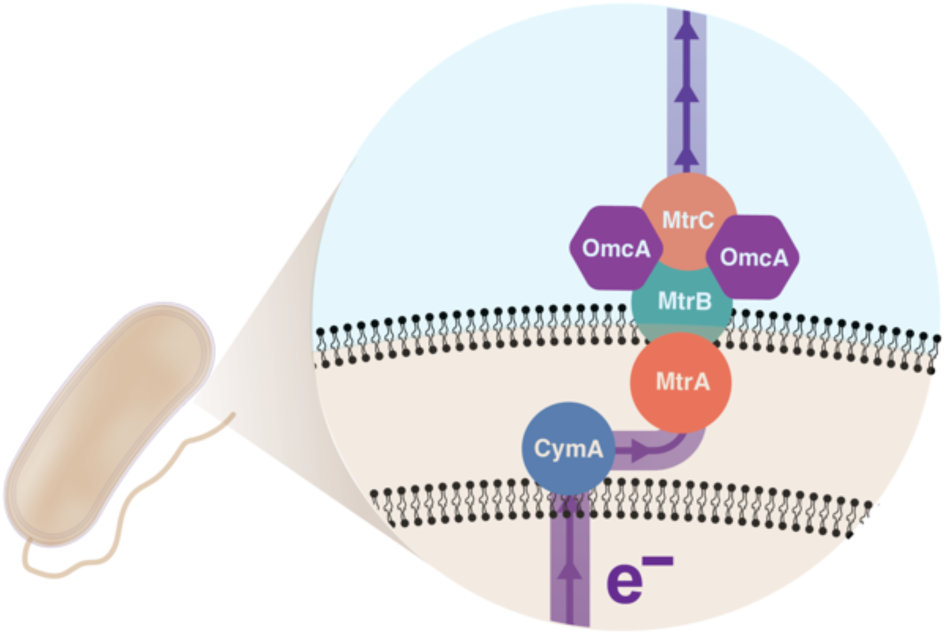
CymA/MtrCAB extracellular electron pathway in *S. oneidensis* MR-1.

While electron transfer rates below wild-type levels are advantageous in certain applications and can be accomplished using genomic deletion strains,^13^ programmable expression of EET proteins remains a non-trivial engineering task. Despite a few examples of inducible EET flux via circuits that control Mtr protein expression,^14–17^ basic engineering parameterizations, such as output/input response curves^18^ (i.e., EET flux vs. inducer concentration) and dynamic range optimization (i.e., induced EET rate/uninduced EET rate), have not been systematically studied. It is not clear the magnitude to which each EET protein affects these parameterizations nor the relative effect of each pathway component. Additionally, many previously engineered circuits exhibited “leaky” expression of EET genes.^17,19^ Finally, most EET genetic circuits have focused on “turning on” EET activity, but “turning off” EET activity at the protein expression level remains largely unexamined.^20^ Taken together, an overall picture of how controlled changes in CymA/Mtr protein expression relate to changes in EET flux is lacking. Developing this understanding is critical for adopting synthetic biology approaches to control EET activity and expanding the versatility of EET in applications such as sensing and catalysis.

To address these challenges, we designed a well-defined and modular expression system for the CymA/Mtr proteins that could be used to rapidly parameterize electroactivity by engineered *S. oneidensis* strains. Transcriptional regulation can be harnessed to precisely control protein expression and design synthetic biology circuits that behave similarly to electronic circuits.^21–23^ By pairing transcriptional regulators (e.g., repressors, activators) with cognate promoter DNA sequences, expression of promoter-downstream genes can be modulated based on the presence/absence of inducing molecules. Using this regulation scheme, transcriptional logic gates can be constructed that facilitate rational programming of complex cellular functions.^24–26^ We hypothesized that rigorous control of EET activity could similarly be enabled using transcriptional logic gates that control expression of single EET genes. By tuning logic gate elements, including RBS sequences and regulatory topology, we predicted that EET activity could exhibit response functions comparable to those observed with fluorescent reporters. Transcriptional control over individual EET components would simplify programming of EET flux and potentially facilitate its integration into previously developed genetic circuits.

In this work, we developed a suite of plasmids to control expression of the EET genes *mtrC, mtrA*, and *cymA. S. oneidensis* strains with appropriate genomic deletions were transformed by these plasmids, and the effects of EET gene transcription and translation were systematically interrogated. When evaluating genetic designs, Fe(III) reduction was used as a high throughput reporter of EET activity. *In situ* reduction kinetics were fitted to transcription-governed models, which allowed us to parameterize strain electron transfer rates and optimize inducible EET turn-on behavior, including dynamic range and fitting to Hill models of gene transcription. After identifying transcriptionally responsive EET constructs, we further demonstrated their utility by constructing NOT gates that functionally turn off EET activity in response to IPTG. Our results show that EET activity can be programmed via expression of single genes and will provide insight on the amenability of EET pathway components for future engineering efforts.

### High Throughput and Real-Time Quantification of Microbial Fe(III) Reduction

To rapidly prototype EET genetic circuits, we developed a simple and high throughput means to detect expression-driven changes in EET flux (Figure 2a). Soluble Fe(III) reduction is a common assay for microbial EET activity and can be measured using the colorimetric dye, ferrozine. Ferrozine complexes microbially generated Fe(II) and has an absorption peak at 562 nm^27^. Inspired by previous *Shewanella* studies^28–30^ and a recent report that analyzed the electroactivity of several microorganisms *en masse*,^31^ we found that adding ferrozine directly to cell suspensions in anaerobically sealed 96-well plates enabled high throughput and real-time monitoring of Fe(III) reduction and EET activity (Supplementary Figure S1). Each experimental 96-plate well was filled with *Shewanella* Basal Medium that contained 1 mg mL^−1^ ferrozine, which showed minimal absorbance change in the absence of bacterial inoculum. In contrast, wells containing ferrozine and *S. oneidensis* cells exhibited color changes attributable to Fe(III) reduction. As plates also contained abiotic Fe(II) standards, this setup enabled rapid detection of Fe(II) levels up to 96 *µ*M across varying biological and chemical conditions. For example, when adding ferrozine to suspensions of *S. oneidensis* MR-1 (wild-type), we observed distinct Fe(III) reduction profiles that follow known *S. oneidensis* metabolisms on different carbon sources.^32^ Bacteria cultured with lactate and pyruvate displayed fast Fe(III) reduction kinetics, relative to starved cultures and those supplemented with metabolically inaccessible acetate (Figure 2b). We also measured drastic kinetic differences between *S. oneidensis* MR-1 and previously characterized Fe(III) reduction-deficient strains.^13^ Cytochrome knockout strains exhibited markedly slower Fe(III) reduction levels, relative to *S. oneidensis* MR-1 (Figure 2c). Together, these results demonstrate that our assay provides a high-throughput means to quantify engineered EET activity via microbial Fe(III) reduction kinetics.

**Figure 2.**
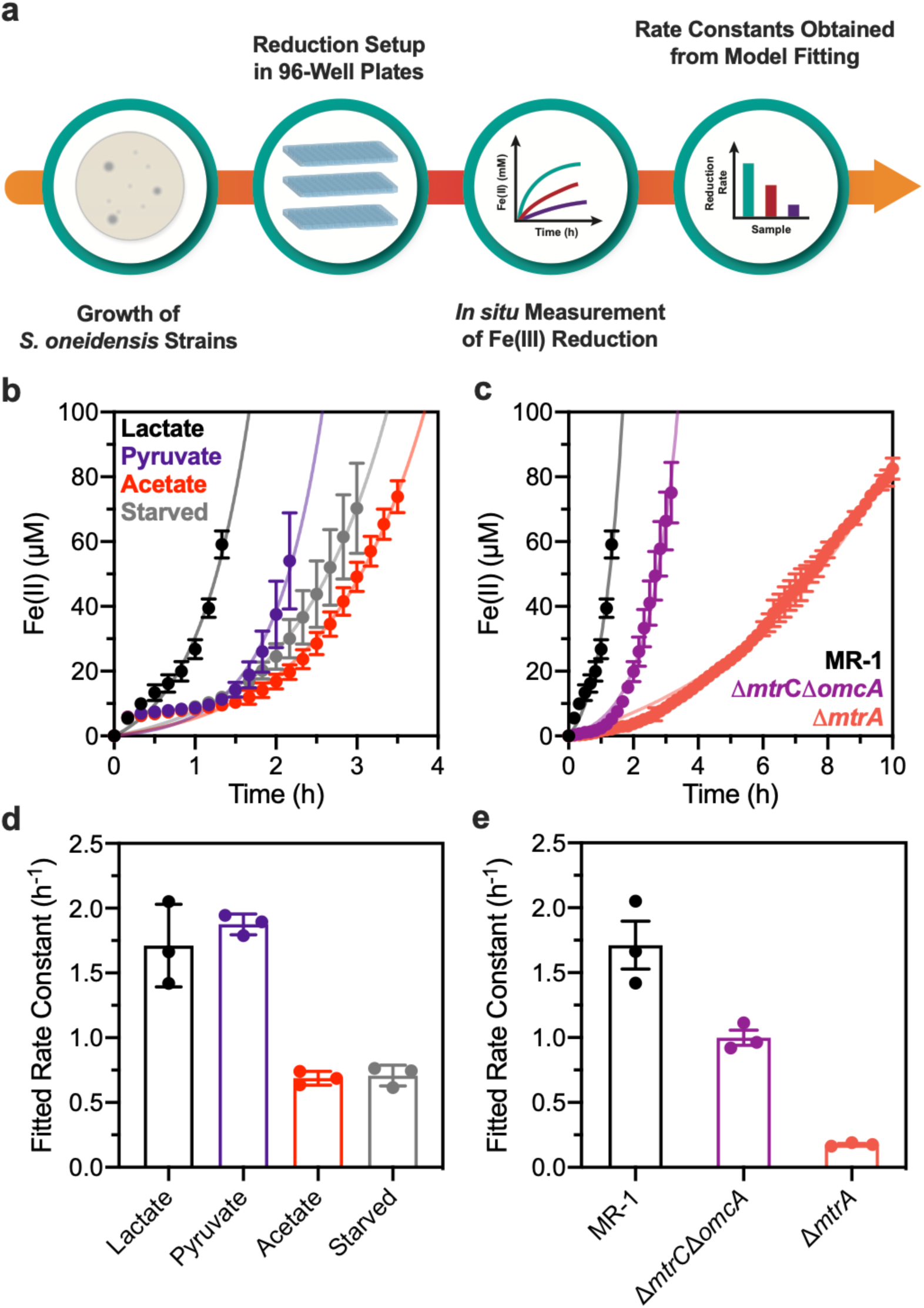
Measurement and modeling of Fe(III) reduction kinetics by *S. oneidensis*. (a) General workflow for 96-well plate Fe(III) reduction setup, kinetic measurements, and obtaining fitted rate constants. (b) Raw Fe(III) reduction kinetics for *S. oneidensis* MR-1 cultured with 1 mg mL^−1^ ferrozine, 5 mM Fe(III) citrate, and either 20 mM sodium lactate (black), 20 mM sodium pyruvate (purple), 20 mM sodium acetate (red) or no carbon source (grey). (c) Raw Fe(III) reduction kinetics for *S. oneidensis* MR-1 (black), Δ*mtrC*Δ*omcA* (purple), and Δ*mtrA* (red) cultured with 1 mg mL^−1^ ferrozine, 5 mM Fe(III) citrate, and 20 mM lactate. Fitted rate constants from the carbon source experiments (b) and different *S. oneidensis* strains (c) are shown in (d) and (e), respectively. Rate constants were obtained by fitting raw kinetic data to a Monod-type model. For (b) and (c), all data represent the mean ± s.d. of raw Fe(II) levels. For (d) and (e), all data represent the mean ± s.e.m of fitted rate constants. All experiments were performed with n = 3 biological replicates.

### A Monod-type Model Connects Steady-State Gene Expression to Fe(III) Reduction

Having established a platform for directly measuring Fe(III) reduction kinetics by various *S. oneidensis* strains and conditions, we next developed a genetic and mathematical framework that could tie EET gene expression to Fe(III) reduction. Despite functional redundancy by other cytochromes and electron transfer proteins,^33^ the CymA/MtrCAB pathway is the primary conduit for electron transfer to soluble Fe(III) by *S. oneidensis*. Knockout strains that lack just one protein in this conduit exhibit significantly reduced rates of Fe(III) reduction relative to wild-type. Given these results, we hypothesized that complementing appropriate knockout strains with a single EET gene from the CymA/MtrCAB conduit would cause Fe(III) reduction kinetics to be rate limited and controllable by expression of that gene. To test this, we designed plasmids that placed *mtrC, mtrA*, and *cymA* downstream of the promoter P_tacsymO_,^34^ which is under control of the plasmid-expressed transcriptional regulator, LacI.^35^ These genes were chosen as they are all cytochromes, span a range of sizes and cellular localizations, and have precedence for altering EET flux via genetic circuits in *S. oneidensis*. Constructed plasmids were used to transform the following knockout strains when appropriate: Δ*mtrC*Δ*omcA*Δ*mtrF*, Δ*mtrA*, Δ*cymA*. These strains exhibit substantially lower Fe(III) reduction capabilities relative to *S. oneidensis* MR-1 that can be restored by complementation.^13,36^ With addition of IPTG and use of various regulator topologies, we varied transcriptional flux for each gene and systematically examined expression effects on Fe(III) reduction rate.

To facilitate comparisons between different tested conditions, we identified cell-intrinsic metrics to analyze our Fe(III) kinetic data. While microbial Fe(III) reduction rates are often normalized by bulk biomass, measurements of cell concentration or protein levels were challenging to perform during *in situ* kinetics. Previous modeling of soluble Fe(III) reduction by *Shewanella* spp. has leveraged Monod-type models to analyze microbial kinetic data^37^. With these exponential models, the fitted exponential constant is a cell-intrinsic, biomass-independent, parameter that determines substrate utilization rates (Supplementary Note S1). Indeed, we found that a Monod-type model fit well to our Fe(II) kinetic data and that qualitative trends in raw Fe(III) reduction kinetics matched fitted rate constants for *S. oneidensis* MR-1 cultured on different carbon sources (Figure 2d) and between different *S. oneidensis* strains (Figure 2e). Using the Monod-type model, extrapolated Fe(II) kinetics from our ferrozine-containing experiments also match previous measurements with these strains in ferrozine-free experiments over longer timescales.^13^ Moreover, culture supplementation with 5 mM Fe(III) citrate was sufficiently high enough such that the fitted rate constant was not appreciably affected by substrate limitation in the range of Fe(II) quantification (0 to 96 *µ*M) (Supplementary Figure S2).

Another benefit of the Monod-type model is that it can directly relate plasmid-based EET gene expression to Fe(III) reduction rate. Similarly to Michaelis-Menten kinetics, the fitted Fe(III) reduction rate constant can be assumed to be proportional to cellular concentration of the rate-limiting protein (i.e., the expressed EET protein) (Supplementary Note S1).^38^ Since EET protein levels are proportional to steady-state mRNA levels during exponential growth,^39^ we predicted that transcriptional logic gate regulation of EET genes would control Fe(III) reduction rates. Thus, the fitted exponential rate constant may serve as a valuable parameter to evaluate how various genetic constructs/experimental conditions affect Fe(III) reduction kinetics.

### RBS Strength Tunes Dynamic Range of Inducible Fe(III) Reduction

Towards designing more complex EET behavior, an important preliminary objective was to determine the dynamic range over which EET gene circuits can turn cellular electron flux “on” relative to an uninduced “off” state. Typically, dynamic range of inducible systems is determined by measuring reporter activity (e.g., GFP fluorescence) under conditions of maximal induction and in the absence of inducer.^40^ These reporter measurements serve as a proxy for the maximum and minimum expression of the reporter gene and can be normalized by cell number or taken at the single cell level. In contrast, our system measures Fe(III) reductase activity in a bulk cell suspension, which makes similar normalizations challenging. Nonetheless, we predicted the fitted reduction rate constants would vary with EET gene expression, and that the ratio of fitted constants could be used to determine the dynamic range of inducible EET circuits.

Maximizing dynamic range in inducible systems is necessary for practical applications, since it affords high signal-to-noise ratios when actuating cellular responses. Thus, we sought to identify inducible EET gene constructs that exhibited high dynamic range, and thus strong control over EET activity. Although dynamic range in our circuit is ultimately determined by changing gene transcription rates, gene translation strength also affects the magnitude of expression levels. For example, even in the absence of inducer, leaky transcription of inducer-regulated genes may occur. If the gene possesses high translational strength, low levels of transcript may be sufficient to saturate protein expression indistinguishably from maximal induction. Indeed, previous LacI regulated *mtrC* constructs we and others have utilized exhibit such behavior.^19,41^ Alternatively, genes with weak translational strengths may exhibit no protein expression under inducing conditions. Since the ribosome binding site (RBS) sequence modulates gene translation rate, this sequence can be optimized for each expressed gene to find a “transcriptional window” that exhibits minimal leaky expression and high dynamic range. Computational tools developed by Salis et al. are frequently utilized to predict and forward engineer gene translation rates.^42^ With their RBSLibrary Calculator, users provide the DNA sequences for a target gene, the desired RBS-upstream region, and the 16S rRNA sequence of the host organism to generate a library of candidate RBS sequences with a spectrum of predicted translational strengths.^43,44^

We leveraged this tool to generate/predict RBS sequences that would optimize the dynamic range of inducible EET gene constructs. For *mtrC, mtrA*, and *cymA*, we assembled a small library of plasmids (5 to 6 members per gene) that varied only in RBS sequence (Figure 3a, Supplementary Table S2). As mentioned above, each plasmid was used to transform an appropriate knockout strain of *S. oneidensis* to avoid native chromosomal expression (Figure 3b-d). To permit insulation from genetic context and potentially enable combination of different EET genes in future assemblies, we also added a self-cleaving ribozyme downstream of the P_tacsymO_ promoter and upstream of each RBS.^45^ Despite the presence of unique ribozymes for each EET gene, the RBSLibrary Calculator includes these different RBS-upstream sequences into translation strength calculations and outputs comparable predictions. We report translation strength for each plasmid as the ΔG_total_, which is the predicted change in Gibbs free energy associated with mRNA binding to the host ribosome relative to the unbound state of both components. Under ideal conditions, the ΔG_total_ for a given gene should be linearly proportional to the logarithm of that gene’s translation rate (and thus steady-state protein levels), with more negative ΔG_total_ values corresponding to higher translation rates.^42^

**Figure 3.**
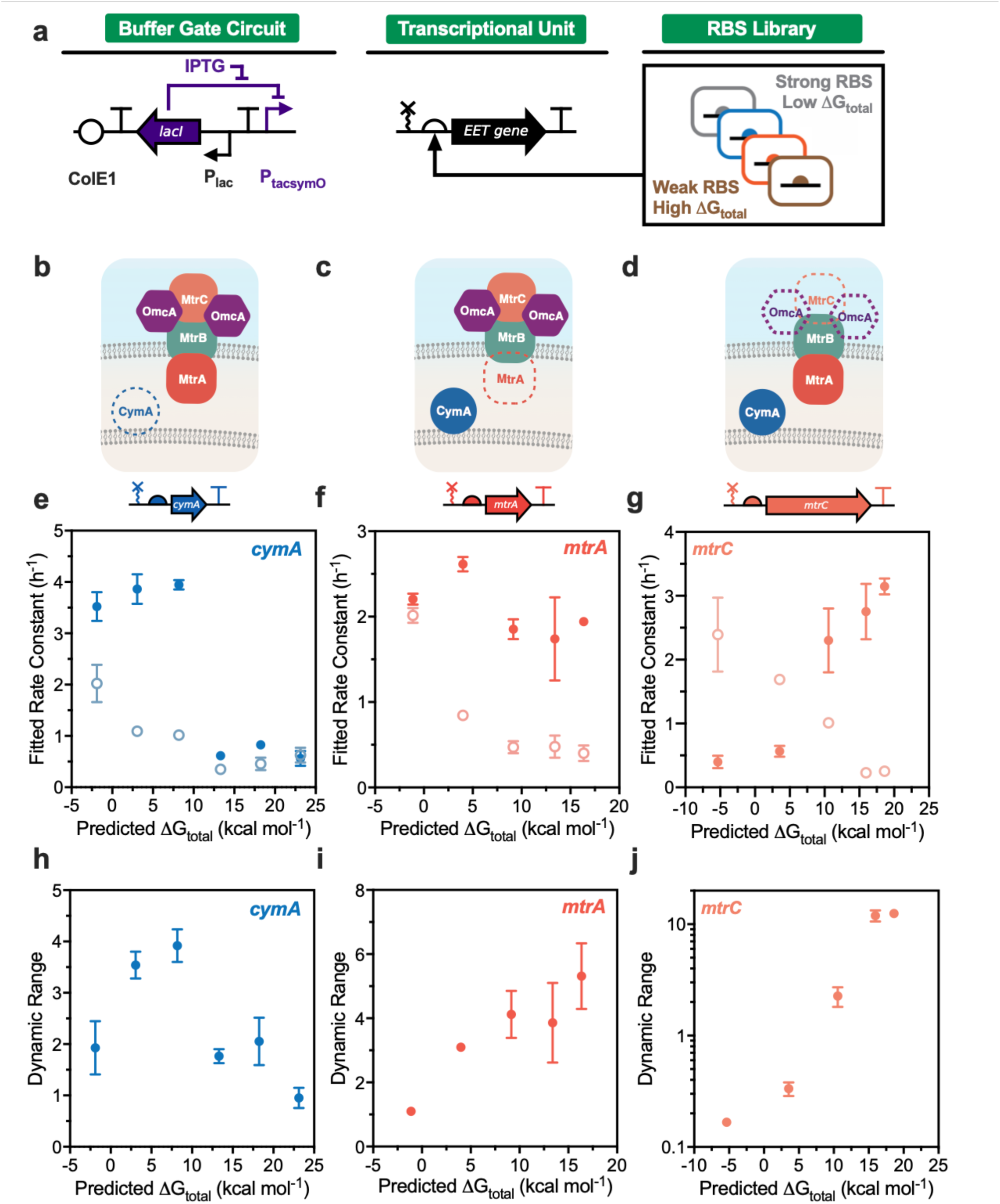
Dynamic range optimization for inducible EET circuits. (a) Diagram of the RBS library/IPTG inducible circuit (Buffer gate) used to control each EET gene transcriptional output. (b) Strain backgrounds and transcriptional unit pairings that were used to assay each RBS library for *cymA* (Δ*cymA*+pCD26r#), (c) *mtrA* (Δ*mtrA*+pCD25r#), and (d) *mtrC* (Δ*mtrC*Δ*omcA*Δ*omcA*+pCD24r#). Dashed outlines represent proteins removed via genomic deletion. Induced (1500 *µ*M IPTG, filled circles) and uninduced (0 *µ*M IPTG, open circles) Fe(III) reduction rate constants for the (e) *cymA*, (f) *mtrA*, and (g) *mtrC* RBS libraries. *cymA* circuit strains were pregrown aerobically and *mtrA*/*mtrC* circuit strains were pregrown anaerobically. For all experiments, IPTG was supplemented immediately prior to measuring Fe(III) reduction kinetics. Rate constants were obtained by fitting raw kinetic data to a Monod-type model. Dynamic range of inducible Fe(III) reduction for the (h) *cymA*, (i) *mtrA*, and (j) *mtrC* RBS libraries. Dynamic range was calculated as the ratio of induced (1500 *µ*M IPTG) and uninduced (0 *µ*M IPTG) fitted rate constants for each replicate. All data represent the mean ± s.e.m. of fitted rate constants or dynamic range for n = 3 biological replicates. Raw data for all replicates is shown in Supplementary Figure S3.

To analyze dynamic range of EET activity for each library member, we measured Fe(III) reduction kinetics in the presence of 0 and 1500 *µ*M IPTG. These conditions facilitate minimal and maximal transcription in our circuit, as determined by sfGFP expression (Figure 4b). To maximize differences in EET protein expression, we chose to induce cells with IPTG immediately prior to Fe(III) reduction. Cells were pregrown uninduced either anaerobically on lactate/fumarate (*mtrC, mtrA*) or aerobically on lactate (*cymA*) for ca. 18 hours and diluted 100-fold into growth medium containing ferrozine and IPTG. Aerobic pregrowth is required for *cymA*-deficient strains due to the protein’s global role in several anaerobic respiratory pathways.^10,36^ After addition of 5 mM Fe(III) citrate, measurement of Fe(III) reduction kinetics began. In the absence of IPTG, we observed that fitted rates constants for all tested genes monotonically increased with more negative ΔG_total_ (Figure 3e-g). This result is consistent with increases in leaky gene expression as translation strength increases and validates the predictive power of the RBSLibrary Calculator in *S. oneidensis*. In contrast, fitted reduction rate constants from IPTG-containing samples showed qualitatively different trends based on the tested gene. With *cymA*, fitted rate constants were comparable for induced and uninduced strains harboring weaker translating library members (Figure 3e). However, rate constants increased to comparable levels for *cymA* strains harboring constructs with predicted ΔG_total_ less than 8.2 kcal mol^−1^. Given the increase in reduction rate constant observed for all uninduced strains with increasing translation strength, a maximum in dynamic range was observed at a relatively intermediate translation strength within our *cymA* library (pCD26r4, ΔG_total_= 8.19 kcal mol^−1^, dynamic range = 3.9 ± 0.3) (Figure 3h). In contrast, maximums in dynamic range were observed for the weakest translating library members of *mtrA* (pCD25r0, ΔG_total_ = 16.4 kcal mol^−1^, dynamic range = 5.3 ± 1.0) and *mtrC* (pCD24r1, ΔG_total_= 18.6 kcal mol^−1^, dynamic range = 12.5 ± 0.3) strains (Figure 3i-j). For induced *mtrA* strains, the fitted rate constants showed minimal change across varying translation strength (Figure 3f). Consequently, dynamic range increased with decreasing translation strength as the background uninduced reduction increases. Interestingly, induced *mtrC* strains exhibited a decrease in fitted rate constant with increasing translation strength, which fell below uninduced levels for constructs with predicted ΔG_total_ less than 5 kcal mol^− 1^ (Figure 3g). This result suggests that *mtrC* expression imposes a metabolic burden at higher translation strengths.^46^ Raw kinetic data from each of the optimal RBS/gene combinations show that different timescales of Fe(III) reduction arise based on construct identity (Supplementary Figure S3). Together, these results highlight the ability of RBS engineering to modulate EET activity and facilitated the identification of three inducible EET plasmids with minimal leakiness and dynamic ranges spanning ca. 4 to 12 in their respective strain backgrounds.

**Figure 4.**
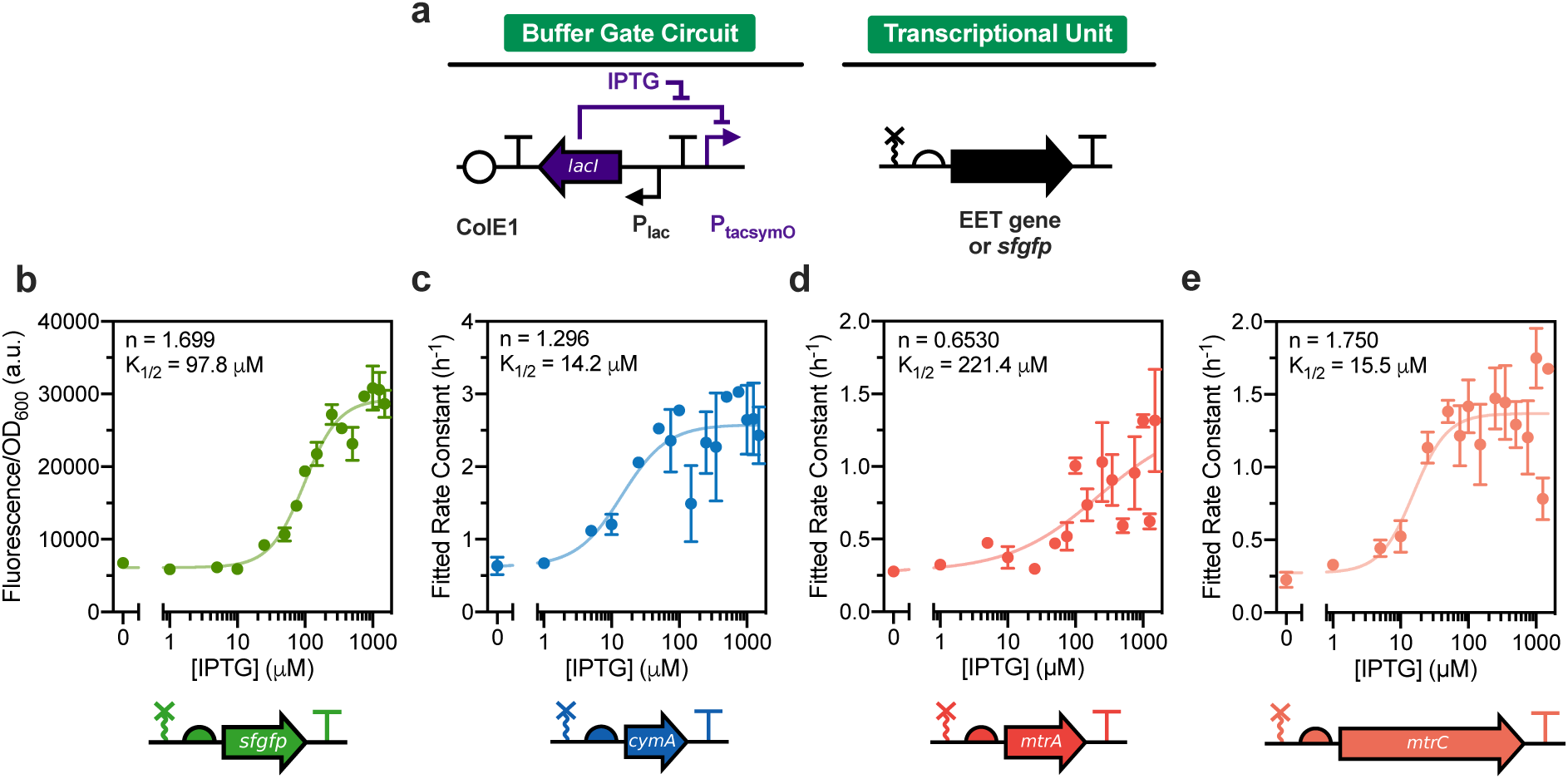
Buffer gate response functions under steady-state EET gene expression. (a) Diagram of Buffer gate circuit used to control each EET gene or *sfgfp* transcriptional output. (b) Fluorescence/OD_600_ response function generated by *S. oniedensis* MR-1 expressing the *sfgfp* Buffer gate circuit. Cells were induced with IPTG for 18 hours during anaerobic growth on lactate/fumarate. To produce mature and functional sfGFP, translation was arrested with 2 mg mL^−1^ kanamycin sulfate and cell aliquots were transferred to an aerobic shaking incubator (30 °C, 250 RPM) for 1 hour prior to fluorescence/optical density measurements. (c) Fe(III) reduction response function generated by aerobically pregrown *S. oneidensis* Δ*cymA* expressing the *cymA* Buffer gate circuit (pCD26r4). (d) Fe(III) reduction response function generated by anaerobically pregrown *S. oneidensis* Δ*mtrA* expressing the *mtrA* Buffer gate circuit (pCD25r0). (e) Fe(III) reduction response function generated by anaerobically pregrown *S. oneidensis* Δ*mtrC*Δ*omcA*Δ*mtrF* expressing the *mtrC* Buffer gate circuit (pCD24r1). For Fe(III) reduction experiments, all strains were pre-induced with IPTG for 18-24 hours prior to measuring Fe(III) reduction kinetics. Cultures were also supplemented with IPTG during kinetic measurements. Rate constants were obtained by fitting raw kinetic data to a Monod-type model. The solid line and Hill parameters shown in each graph were obtained by fitting individual replicates to a four-parameter activating Hill function with weighted error. To improve Hill function fitting for (d), the minimum and maximum Hill function values were constrained by the mean fitted rate constants at 0 and 1500 *µ*M IPTG, respectively. All data represent the mean ± s.e.m. of fitted rate constants for n = 3 biological replicates. Raw data for all replicates is shown in Supplementary Figure S12.

### Inducible Fe(III) Reduction Occurs in the Absence of Ferrozine

To confirm that inducibility was not an artifact of our real-time kinetics setup, we also performed inducible expression for each optimized EET gene in lactate/Fe(III) citrate medium that lacked ferrozine. Indeed, we observed that Fe(III) reduction remained highly inducible via IPTG for each construct (Supplementary Figure S4). In these experiments, we concomitantly measured viable cell density and observed comparable, albeit slightly higher, growth yields for induced samples relative to uninduced. Although growth dynamics were not continuously monitored, cell concentrations for each strain differed based on the presence of IPTG by at most ca. 2-fold after 8 hours, while Fe(II) levels varied 7-to 65-fold. This result suggests that relative differences in biomass do not appreciably contribute to observed differences in Fe(III) reduction rate between induced/uninduced samples and that inducible Fe(III) reduction is primarily controlled by EET protein expression. Additionally, our viability measurements show that inducibility is not due to spurious effects from cell lysis or plasmid maintenance. We also examined the early timescale of inducible Fe(III) reduction for the optimized *mtrC* construct in the absence of ferrozine. Similar to ferrozine-containing experiments, Fe(II) concentration increased in the presence of IPTG after ca. 2-3 hours, while uninduced samples remained at baseline levels (Supplementary Figure S5). Collectively, these results demonstrate that ferrozine does not markedly alter inducible Fe(III) reduction and further validate our real-time methodology as a means to assay EET gene expression.

### Reduction Rate Response Functions from Steady-State EET Gene Expression

Using our translation-optimized EET gene expression plasmids, we next examined how Fe(III) reduction kinetics changed with varying transcription rate. Since our RBS library screening aided in identifying RBS/EET gene pairs that display high dynamic range between maximal (1500 *µ*M IPTG) and minimal (0 *µ*M IPTG) induction, we hypothesized that graded EET activity would be measured when testing intermediate inducer concentrations. Given that EET activity in our system should be proportional to steady-state mRNA levels of the expressed EET gene and transcriptionally regulated via LacI (Figure 4a), we further reasoned that generated response functions (i.e., fitted rate constant vs. IPTG concentration) may exhibit Hill function-type responses to changing IPTG concentration and functionally behave as Buffer logic gates.

To promote steady-state behavior and minimize simultaneous effects of growth and dynamic protein expression, we initially pre-induced each EET construct at various IPTG concentrations for 24-hours in lactate/fumarate-containing growth medium. Subsequently, these cultures were diluted 100-fold into growth medium containing the same concentrations of IPTG and ferrozine, and Fe(III)-citrate was finally added prior to kinetic measurements. As predicted, we observed that fitted rate constants increased for all tested EET genes with increasing concentration of IPTG (Figure 4c-e). Generally, each response function followed a sigmoidal shape and could be modeled using an activating Hill function. Similar fitted cooperativity (n = 1.296 and 1.750) and threshold-response (K_1/2_ = 14.2 and 15.5 *µ*M IPTG) values were observed between *cymA* and *mtrC* (Figure 4c,e). These fitted Hill parameters also comparable favorably to those generated by a fluorescent *sfgfp* circuit that was anaerobically induced with IPTG under steady-state expression (n = 1.699, K_1/2_ = 97.8 *µ*M IPTG) (Figure 4b). This is expected if transcription of each gene is controlled by the same regulator (LacI) and promoter (P_tacsymO_) and was evidence that gene transcription rate limits EET activity in our constructs. While the *mtrA* construct exhibited a more gradual increase in Fe(III) reduction rate (n = 0.6530, K_1/2_ = 221.4 *µ*M IPTG), it still demonstrated an ability to turn-on EET activity in response to IPTG (Figure 4d). Given that empty vector controls showed baseline or lower than uninduced rates of Fe(III) reduction (Supplementary Figure S6), increases in EET activity appear to be due to IPTG-driven increases in EET gene expression for all Buffer gate constructs.

To verify if differential EET activity arose from transcriptional control of EET protein levels, we analyzed whole cell lysate from each EET gene expression construct via SDS-PAGE and heme-staining. For the *mtrC* construct, we observed high molecular weights bands (ca. 250 kDa) that increased in intensity with increasing concentration of supplemented IPTG (Supplementary Figure S7a). Bands in this size range have previously been associated with MtrC-containing complexes.^47^ For the *cymA* construct a faint but noticeable band migrated around the size of CymA (19 kDa) (Supplementary Figure S7c).^48^ While IPTG-dependent differences could not be discerned by the naked eye, densitometry analysis indicated that band intensity increased with IPTG concentration above uninduced levels (Supplementary Figure S7d). Although we could not identify bands whose formation/disappearance was a result of MtrA expression (Supplementary Figure S7b), the MtrC/CymA heme staining results are evidence that functional cytochromes can be generated from our expression system. Together, this data demonstrates that transcriptionally regulated *cymA, mtrA, and mtrC* constructs can controllably turn on EET activity (i.e., Buffer gate behavior) and suggests that changes in protein expression are linked to changing Fe(III) reduction rates.

### A Polynomial Model Describes Dynamic Gene Expression and Fe(III) Reduction

While these previous experiments demonstrated that our inducible constructs afforded predictable control over EET activity under conditions closest to steady-state protein expression, there are many applications where dynamic control of protein expression and EET activity would be advantageous (e.g., sensing, catalysis).^15,41^ Thus, we also examined EET response functions by the *cymA, mtrA*, and *mtrC* constructs in response to non-steady state gene expression. As described above, we previously fit Fe(III) reduction kinetics to a Monod model of substrate utilization. However, a major assumption of Monod models is that transcriptomic and proteomic changes during growth do not affect the fitted rate constant.^49^ In practice, this means that engineered changes to mRNA and protein levels should rapidly reach steady-state. As this may not be the case under certain experimental conditions, we also examined a model that accounted for the dynamics of protein expression immediately after induction. Under the assumption of constant biomass (i.e., minimal growth) and rapid equilibrium of steady-state mRNA levels, a simplified dynamic induction model predicts a linear increase in the cellular concentration of expressed EET proteins with time (Supplementary Note S2). The minimal growth assumption appears valid based on comparable cell counts measured for each induced/uninduced EET construct during dynamic expression (Supplementary Figure S4). While this model exhibits an accelerating Fe(III) reduction rate similar to Monod-based exponential models, kinetics are more accurately described by a second-order polynomial. Using this polynomial model, the fitted constant preceding the time-squared term is also predicted to be proportional to steady-state mRNA levels and should be controllable in our system. As our system is likely somewhere in-between the limits of no-growth/dynamic protein levels and cell growth/steady-state protein levels, we expected that both models could aid characterization of dynamic EET gene expression and Fe(III) reduction.

### Dynamic Expression of EET Genes Predictably Controls Reduction Rate

To evaluate dynamic expression of EET genes, cells were pregrown in the absence of inducer and induced with IPTG immediately prior to Fe(III) citrate addition. Initially, we mimicked our steady-state experiments by preculturing the *mtrA* and *mtrC* constructs anaerobically and the *cymA* construct aerobically. Using the polynomial model to fit Fe(III) reduction kinetics, the response function generated by the *cymA* construct fit strongly to an activating Hill equation (n = 1.176, K_1/2_ = 79.54 *µ*M) (Figure 5b) and fitted Hill parameters were comparable to the response function generated via steady-state *cymA* expression (Figure 4c). Rate constants obtained from the Monod-type model yielded similar *cymA* response function behavior (Supplementary Figure S8f). In contrast, response functions generated by the *mtrA* and *mtrC* strains fit poorly to Hill models and exhibited low Hill coefficients (n = 0.5492 to 0.7049) for both polynomial and Monod-type model fitting (Supplementary Figure S8b-e). While polynomial model fitting produced response functions with higher Hill coefficients relative to the Monod-type model, coefficient values for each construct were still substantially lower than what was expected based on the *cymA* and sfGFP circuits (Figure 4b). While the *mtrC* construct exhibited a steep EET turn-on under steady-state expression, turn-on behavior under dynamic expression was substantially more graded. The slope of the *mtrA* dynamic expression response function was similar to its steady-state expression counterpart and to the *mtrC* dynamic expression response function. Together, this was suggestive that other cellular processes (e.g., protein maturation/localization, background Fe(III) reduction) decrease sensitivity of the EET output to IPTG during dynamic induction.

**Figure 5.**
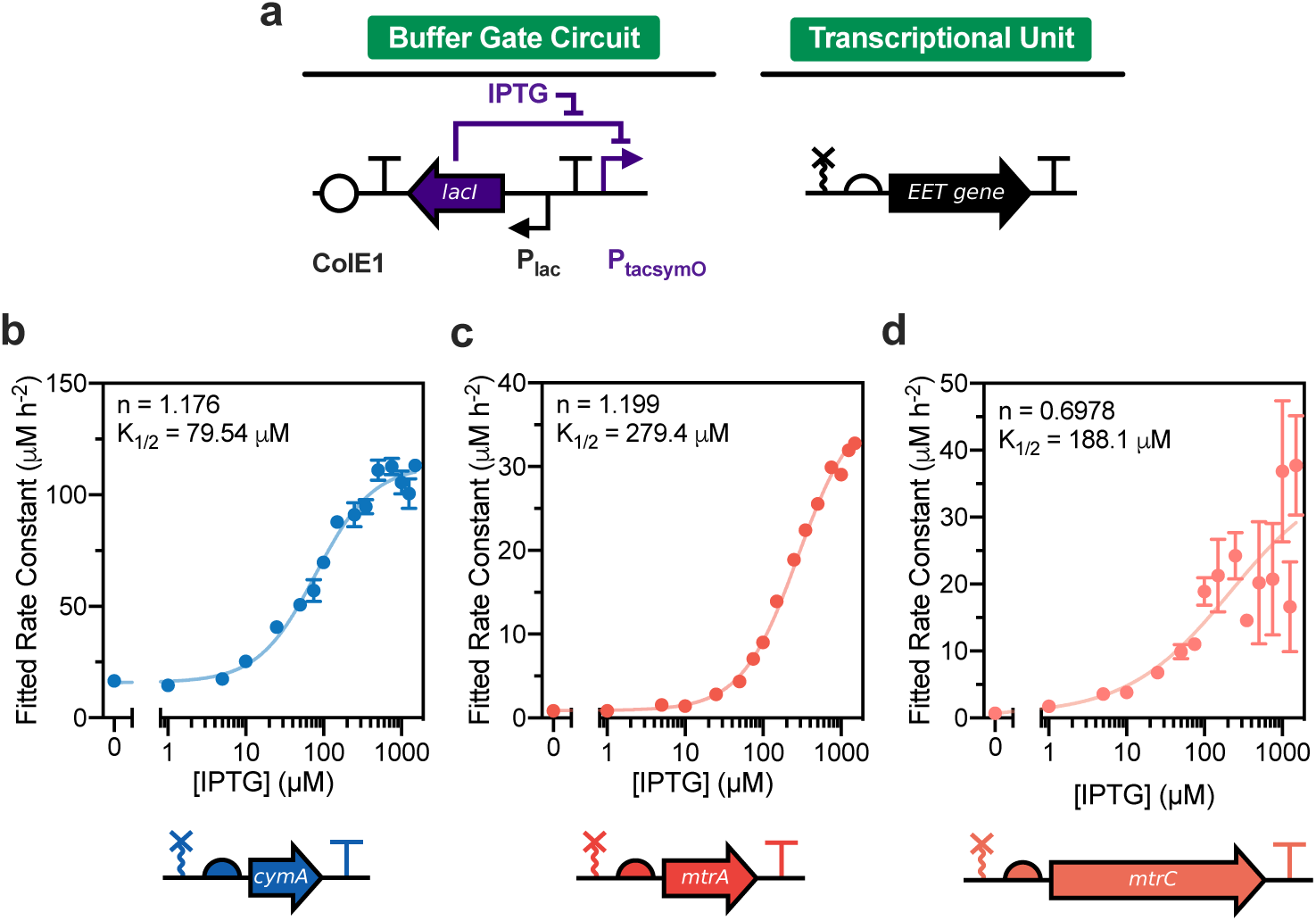
Buffer gate response functions under dynamic EET gene expression and from aerobic pregrowth. (a) Diagram of Buffer gate circuit used to control each EET gene transcriptional output. (b) Fe(III) reduction response function generated by *S. oneidensis* Δ*cymA* expressing the *cymA* Buffer gate circuit (pCD26r4). (c) Fe(III) reduction response function generated by *S. oneidensis* Δ*mtrA* expressing the *mtrA* Buffer gate circuit (pCD25r0). (d) Fe(III) reduction response function generated by *S. oneidensis* Δ*mtrC*Δ*omcA*Δ*mtrF* expressing the *mtrC* Buffer gate circuit (pCD24r1). In these experiments, all strains were pregrown aerobically and IPTG was supplemented immediately prior to measuring Fe(III) reduction kinetics. Rate constants were obtained by fitting raw kinetic data to a polynomial model. The solid line and Hill parameters shown in each graph were obtained by fitting individual replicates to a four-parameter activating Hill function with weighted error. All data represent the mean ± s.e.m. of fitted rate constants for n = 3 biological replicates.

Encouraged by the predictability of the *cymA* response function, we hypothesized that aerobic pregrowth conditions may facilitate higher sensitivity of EET flux to IPTG. Most of the EET machinery is downregulated under aerobic conditions, which we predicted could minimize background Fe(III) reduction.^50,51^ Furthermore, dynamic expression in parallel with aerobic to anaerobic transition may ensure that all EET pathways, including those under exogenous control, turn on simultaneously. Thus, we examined dynamic expression by the *mtrA* and *mtrC* constructs after aerobic pregrowth. Excitingly, we found that response functions for the *mtrA* construct fit more strongly to activating Hill models (Figure 5c). Kinetic fitting using both the polynomial and Monod-type models generated similar values for fitted Hill parameters (Figure S8g). Additionally, the fitted Hill coefficients for the *mtrA* construct were comparable to those for *cymA* and *sfgfp* circuits and overall kinetic variability significantly decreased. In contrast, the *mtrC* construct only showed a marginal improvement in fitting to all models (Figure 5d, Supplementary Figure S8h). These collective results demonstrate that our engineered *cymA, mtrA*, and *mtrC* constructs can controllably turn on EET activity in response to both steady-state and dynamic transcriptional cues.

### NOT Gate Circuits Turn Off Fe(III) Reduction in the Presence of IPTG

Having developed a set of EET gene expression constructs that turn on EET activity via transcriptional regulation, we next asked if similar genetic designs could be leveraged to turn off cellular electron flux. The NOT gate is the simplest expression-based logic gate that inverts the signal from an inducer molecule to turn off expression of a target gene.^22,52^ Mechanistically, an inducer molecule (e.g., IPTG) stimulates expression of a repressor protein, which subsequently downregulates the target gene. Since our *cymA* and *mtrC* constructs were optimized to exhibit maximal transcriptional responsiveness with high dynamic range, we hypothesized that converting their Buffer gate constructs into NOT gate designs would enable transcriptional downregulation of these genes. Accordingly, NOT gate genetic architectures should facilitate EET turn-off behavior in the presence of IPTG.

We designed NOT gate plasmids for the *cymA* and *mtrC* constructs, using TetR as the repressor that regulates EET gene transcription (Figure 6a).^53^ NOT gate transcriptional outputs were identical to our optimized Buffer gate designs for *cymA* and *mtrC* (i.e., ribozyme-optimized RBS-target gene-terminator), but instead of regulation via P_tacsymO_, each transcriptional unit was placed downstream of the promoter P_tet_. Genes under control of P_tet_ are constitutively expressed in the absence of the TetR and downregulated in its presence. In our NOT gate plasmids, TetR expression was controlled via LacI/P_tacsymO_. Thus, IPTG should elevate TetR levels and subsequently decrease EET gene expression in appropriate knockout strains (Δ*mtrC*Δ*omcA*Δ*mtrF* and Δ*cymA*) transformed by these plasmids. Although EET activity can readily be turned on through dynamic expression of EET genes, turning off EET activity via transcriptional regulation is more challenging. Using NOT gate architectures, regulated EET genes are maximally expressed in the absence of inducer. Upon adding IPTG, not only must downregulation of EET gene transcription be considered, but also the fate of preexisting EET proteins. While EET proteins can potentially be diluted through cell division,^54^ this process may occur at timescales slower than what is required for functional NOT gate behavior.

**Figure 6.**
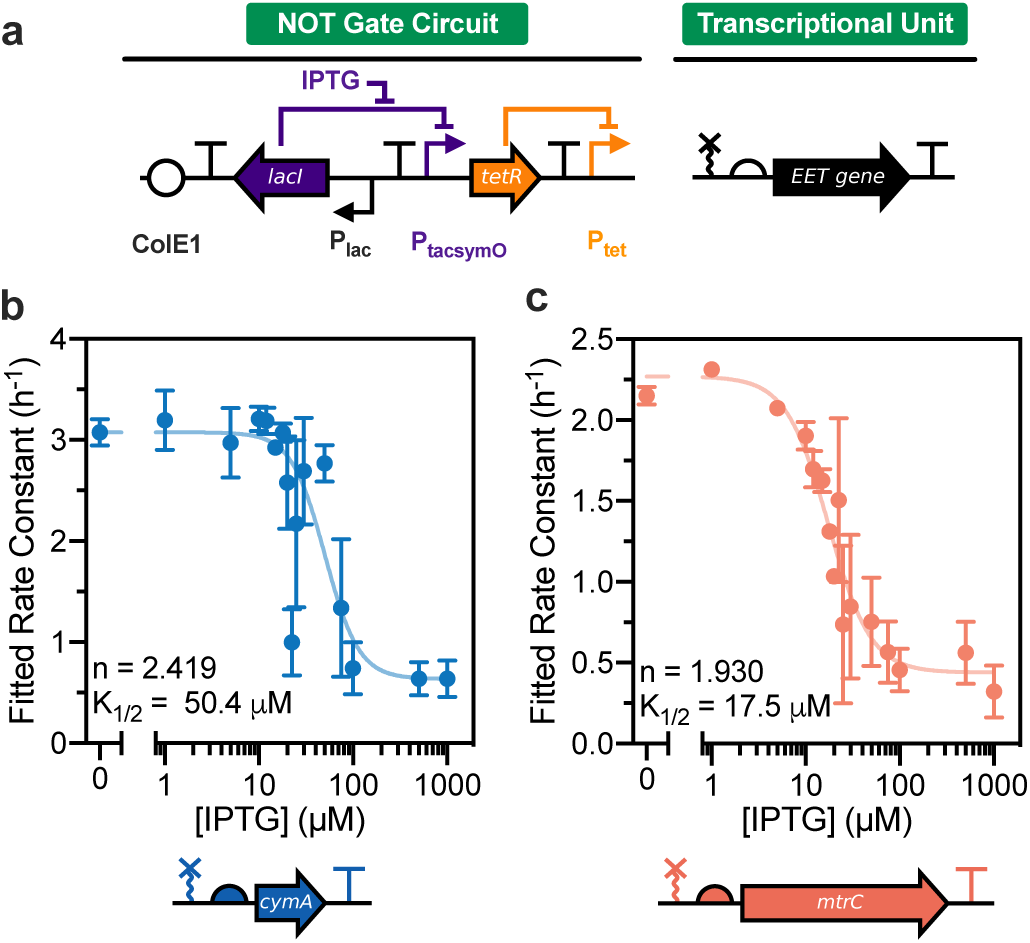
NOT gate response functions under steady-state EET gene expression. (a) Diagram of NOT gate circuit used to control each EET gene transcriptional output. (b) Fe(III) reduction response function generated by aerobically pregrown *S. oneidensis* Δ*cymA* expressing the *cymA* NOT gate circuit (pCDd3r4). (c) Fe(III) reduction response function generated by anerobically pregrown *S. oneidensis* Δ*mtrC*Δ*omcA*Δ*mtrF* expressing the *mtrC* NOT gate circuit (pCDd1). In these experiments, all strains were pre-induced with IPTG for 18-24 hours prior to measuring Fe(III) reduction kinetics. Cultures were also supplemented with IPTG during kinetic measurements. Rate constants were obtained by fitting raw kinetic data to a Monod-type model. The solid line and Hill parameters shown in each graph were obtained by fitting individual replicates to a four-parameter deactivating Hill function with weighted error. To improve Hill function fitting for (b), the maximum and minimum Hill function values were constrained by the mean fitted rate constants at 0 and 1500 *µ*M IPTG, respectively. All data represent the mean ± s.e.m. of fitted rate constants for n = 3 biological replicates.

To avoid complications from dynamic expression/induction and achieve steady-state expression levels, an 18-24 hour IPTG pre-induction was performed for each NOT gate prior to Fe(III) reduction. Response functions were generated using a range of IPTG concentrations and we found that fitted rate constants were reasonably approximated by deactivating Hill functions for both the *cymA* (n = 2.419; K_1/2_ = 50.4 *µ*M) and *mtrC* (n = 1.930; K_1/2_ = 17.5 *µ*M) constructs (Figure 6b-c). In the absence of IPTG, both constructs exhibited rapid Fe(III) reduction and fitted rate constants were similar to those observed with our maximally induced EET turn-on circuits. This suggested that both *mtrC* and *cymA* were being constitutively expressed as expected from the NOT gate design. In contrast, cells preinduced with high levels of IPTG exhibited slower Fe(III) reduction kinetics, with fitted rate constants comparable to both uninduced *mtrC* and *cymA* Buffer gate constructs. To control for the effect of TetR expression on Fe(III) reduction, we also constructed an empty NOT gate plasmid that increased TetR expression in response to IPTG, but lacked an EET gene downstream of the P_tet_ promoter. All knockout strains transformed by this plasmid exhibited low levels of Fe(III) reduction and minimal change in reduction rate with IPTG addition (Supplementary Figure S9). While, *S. oneidensis* MR-1 transformed by this plasmid exhibited a slight drop in Fe(III) reduction in the presence of IPTG, induced and uninduced rate constants remained substantially higher than empty vector or NOT gate transformed knockout strains. This further evidenced that changes in Fe(III) reduction were due to NOT gate-controlled EET gene expression and not spurious effects from varying TetR levels.

We briefly examined dynamic expression experiments with the *mtrC* NOT gate. As predicted, only a marginal decrease in Fe(III) reduction rate was observed when IPTG was added immediately prior to kinetic measurements (ca. 35%) relative to decreases observed with steady-state expression (ca. 85%) (Supplementary Figure S10, Figure 6c). Attempts to accelerate growth-based protein turnover with addition of fumarate were similarly hampered by poor EET turn-off behavior, relative to steady-state experiments (Supplementary Figure S10). Although dynamic expression will require further optimization, our results indicate that it is possible to program EET turn-off by downregulating expression of EET genes.

## Discussion

Developing rigorous and predictable control over EET flux is of critical importance to expand the utility of electroactive microbes in redox-driven technologies.^11,55^ In this work, we asked if expression of single EET genes (*cymA, mtrA, mtrC*) in *S. oneidensis* knockout strains could enable such control. By making simple and systematic changes to translation and transcription of these genes, we explored the design space of programmable Fe(III) reduction kinetics.

Although there are notable examples of engineered genetic circuits that control EET activity in *S. oneidensis*, many of these circuits simultaneously regulate multiple genes (e.g., *mtrCAB* operon,^17,20^ NADH/flavin metabolite pathways^12,56^) or lack detailed engineering parameterizations typically measured for fluorescence-parameterized circuits (e.g., response functions, dynamic range optimization). Although more systematic optimizations have been performed to incorporate the Mtr proteins into *Escherichia coli*, this organism requires extensive engineering to facilitate proper functioning of pathway components.^19,46,57^ Since the underpinnings of EET performance can be challenging to interpret, we adopted a reductionist approach by studying single EET genes and their contribution to EET flux. Expressing each gene in appropriate *S. oneidensis* knockout strains allowed us to abrogate native expression, functionally generate each EET protein and associated conduit proteins, and isolate each gene’s influence on EET parameters. Moreover, we chose to target the CymA/Mtr conduit since these proteins are somewhat orthogonal to overall cellular metabolism, relative to more universal metabolites (e.g. NADH, flavins, quinones). In contrast to electrode-based setups, Fe(III) reduction provided a convenient and high throughput assay to analyze EET activity. Notably, our results demonstrate that fitted Fe(III) reduction rate constants from engineered strains follow predictable behavior, akin to circuits where fluorescent reporters are the output.^22^ While cellular physiology likely differs between knockout strains (e.g., loss of one cytochrome may upregulate expression/maturation of others)^46^ and absolute kinetics are subject to change based on plasmid composition (e.g., origin of replication, resistance marker),^34,58^ our system could identify pairings of engineered plasmids and genetic backgrounds that yielded the desired circuit function.

For example, a major challenge in the electromicrobiology community is engineering of EET circuits that exhibit minimal leakiness and high signal-to-noise outputs.^11^ More broadly, developing genetic circuits with these properties is a ubiquitous goal in synthetic biology.^18,22^ With other engineered genetic circuitries, RBS optimization for target genes is typically performed to address this.^53^ Thus, we developed a small library of plasmids for each tested EET gene to first identify the range of inducibility afforded by different RBS sequences/translation strengths. After analyzing Fe(III) reduction kinetics by all library members, we pinpointed RBS sequences for each EET gene that facilitated the highest dynamic range in EET activity. Notably, the optimal *mtrC* and *mtrA* plasmids possessed the weakest translation strength in their respective libraries (Figure 3f-g). Like other outer membrane proteins, MtrC has a low turnover rate (half-life ca. 16 hours) relative to intracellular proteins.^59^ MtrC also forms a large multi-protein complex (MtrCAB) that may further stabilize it from degradation.^60,61^ Moreover, we observed that dynamic range of *mtrC* constructs decreased below 1 with increasing translation strength (Figure 3j), which suggested that high levels of MtrC expression impose a metabolic burden.^46^ Taken together, these factors may explain why only the weakest translation strength construct facilitated high dynamic range. While the degradation rate of MtrA has not been studied, it is also part of the MtrCAB complex, and thus its RBS library likely exhibits similar trends to *mtrC* due to low protein turnover.^62^ Conversely, the small size of CymA (19 kDa) relative to MtrC (76 kDa) and MtrA (35 kDa), along with its more intracellular localization, may increase its susceptibility to proteolytic degradation and result in a higher turnover rate. This would also explain why the optimal *cymA* construct possessed a more intermediate translation strength (Figure 3e). Overall, we identified transcriptional units (ribozyme, RBS, and terminator sequences) for *cymA, mtrA*, and *mtrC* that exhibited a spectrum of dynamic ranges, which could inform the choice of EET construct based on application needs.

We leveraged RBS optimization of each EET gene to forward engineer EET turn-on and turn-off behavior in response to logic gate-driven transcriptional cues. While there are several means to regulate the expression and activity of electron transfer proteins within cells,^20,63^ a benefit of transcriptional logic gates is that desired gene expression behavior is relatively straightforward to program. Using the two simplest logic gates, we validated our underlying hypothesis that EET flux can be precisely controlled via transcriptional modulation of single EET genes. Upon addition of IPTG to each Buffer gate strain, increases in EET protein levels were concomitant with increases in Fe(IIII) reduction rate (Figures 4-5). Although direct measurement of MtrA protein expression posed challenges, our heme staining results suggest that MtrC and CymA are being functionally expressed (Supplementary Figure S7). We also note that all of our circuit plasmids rescued Fe(III) reduction deficiencies that have been previously characterized via knockout complementation.^13,36^ Additionally, minimal changes in cell growth were observed between induced/uninduced strains, further suggesting that kinetic differences were a result of protein expression (Supplementary Figure S4). Furthermore, we observed that fitted Fe(III) reduction rate constants vs. inducer IPTG concentrations (i.e., EET response functions) fit well to Hill models for both our Buffer and NOT gate architectures (Figures 4-6). Since fluorescent protein-based response functions can be readily parameterized by such models (Figure 4b),^18^ these collective results support a scheme whereby transcriptional regulation controls EET protein levels and thus EET flux. Excitingly, our NOT gate circuits were able to dial down EET activity, albeit only at steady-state protein expression levels. As shown from our experiments, utilizing NOT gate architectures under dynamic conditions remains an outstanding challenge and is likely due to limited turnover of the expressed EET proteins (Supplementary Figure S10). Since growth-based dilution is the primary contributor to protein level decreases,^54^ future studies can potentially optimize bacterial respiration onto secondary substrates to improve such decreases for target EET proteins. Altogether, our system demonstrates that EET genes may serve as useful outputs for other transcriptionally controlled circuits, such as two-component sensors and stimuli-responsive logical operators.^26,64^

As determined by the value of fitted Hill coefficients, the sensitivity of EET turn-on activity to IPTG appears to strongly depend on the identity of the expressed EET gene/genetic background and strain pregrowth conditions. As our sfGFP Buffer gate and EET turn on circuits utilize the same transcriptional regulatory mechanism (Figure 4a), fitted Hill coefficients should be comparable for both systems if transcriptional control rate-limits functional activity. Hill parameters for EET response functions that deviate from those measured via sfGFP suggests that other cellular processes compete with transcriptional control.^18,46^ For example, *cymA* and *mtrC* strains that were preinduced with IPTG exhibited Hill coefficient values closest to those determined for sfGFP (n = 1-2) (Figure 4b-c,e), suggesting strong transcriptional control is facilitated via steady-state protein expression. For dynamic expression experiments, we observed lower than expected Hill coefficients (n < 1) for anaerobically pregrown *mtrC* and *mtrA* strains (Supplementary Figure S8). This lower sensitivity to IPTG is reasonable given that these genes encode proteins with numerous post-translational processing steps (e.g., maturation, periplasmic/extracellular localization) that may rate-limit their functional activity when dynamically expressed.^46^ Moreover, other cytochromes may be expressed during anaerobic pregrowth that contribute to background Fe(III) reduction.^13^

Interestingly, we observed that some aerobically pregrown strains (*cymA* and *mtrA*) exhibited a more characteristically sensitive response to IPTG during dynamic expression and fit better to Hill models relative to anaerobic pregrowth (Figure 5b-c). Given that relative growth rates for these strains were not significantly different based on the presence/absence of IPTG (Supplementary Figure S4), we primarily attribute this sensitivity to a decrease in background reduction from spurious EET pathway expression. Under aerobic conditions, much of the EET machinery in downregulated in *S. oneidensis*.^50,51^ Even when switched to anaerobic conditions, Fe(III) reduction capabilities by aerobically pregrown *S. oneidensis* are considerably diminished relative to anaerobic pregrowth.^65^ Thus, the switch to anaerobic Fe(III) respiring conditions likely causes reduction by aerobically pregrown strains to be more rate-limited by plasmid-based EET gene expression. For example, the *mtrA* construct’s genetic background (Δ*mtrA)* has functional redundancy by other cytochromes and is less “insulatory” than the other tested strains (Δ*mtrC*Δ*omcA*Δ*mtrF* and Δ*cymA*),^13,33^ which may explain why the aerobic pregrowth facilitated markedly higher IPTG sensitivity (Figure 5c). Future engineering efforts may seek to identify other strain backgrounds that improve transcriptional control of EET activity. Collectively, our results demonstrate that inducible EET activity can follow a spectrum of sensitivity to inducer stimuli, which may be advantageous based on application needs such as switch-like behavior for computation or graded responses for sensing/detection.^15,66,67^

Logic gate formalisms were originally intended to abstract electrical circuit construction and aid the development of more complex electronic devices. Synthetic biologists have adapted these concepts to generate countless transcriptional circuits,^68^ which range from simple Buffer gates^35^ (a single inducer causes gene expression “turn on”) to complex multi-layered logic constructs,^40,66,67,69,70^ where one regulator modulates the expression of multiple regulators and actuators. In this work, we showed that transcriptional logic gate regulation can be extended to EET protein expression and can consequently control electron transfer to soluble Fe(III). Notably, simple exponential and polynomial relationships captured the dynamics of Fe(III) reduction. From these models, we demonstrated that EET protein actuations can be parameterized similarly to other transcriptionally controlled logic gates (i.e., fitting rate constants to Hill models) and rationally programmed to exhibit desired behavior. Although we only examined the two simplest transcriptional logic gates, the ability to turn on and turn off EET flux is foundational for constructing more complex circuit architectures. Given the respiratory flexibility of the CymA/Mtr pathway, our system is potentially extendable to other metals and EET acceptors. We expect that this work will serve as the basis for future engineering of logic gate-driven EET activity and further blend the fields of synthetic biology and electrical engineering.

## Author Contributions

C.M.D. and B.K.K. conceived the project, analyzed the results, and wrote the manuscript; C.M.D. designed the genetic constructs and performed the experiments.

## Notes

The authors declare no competing financial interest.

## Acknowledgements

This work was supported by the Welch Foundation (Grant No. F-1929) and by the National Institute of General Medical Sciences of the National Institutes of Health under Award Number R35GM133640. The content is solely the responsibility of the authors and does not necessarily represent the official views of the National Institutes of Health Additional research support was provided by the National Science Foundation through the Center for Dynamics and Control of Materials: an NSF Materials Research Science and Engineering Center under Cooperative Agreement DMR-1720595. We would like to thank Prof. Jeffrey Gralnick (U. Minnesota) for generously providing *S. oneidensis* strains Δ*mtrC*Δ*omcA*Δ*mtrF* and Δ*cymA*. We also acknowledge the ICMB Core Facilities at UT Austin for performing DNA sequencing.

## Materials and Methods

### DNA and Strain Construction

All bacterial strains, plasmids, geneti circuit maps, and sequence information for each genetic part are detailed in the Supplementary Information. All plasmids were assembled via Golden Gate cloning procedures using enzymes (BsaI, SapI, BsmBI) and buffers from New England Biolabs. DNA fragments used in Golden Gate cloning^71^ were generated via partial/whole-plasmid PCR (Phusion High Fidelity DNA Polymerase, New England Biolabs) or commercially synthesized (gBlocks, Integrated DNA Technologies). Generally, 10 *µ*L Golden Gate reactions were set up that contained 10 fmol of pCD plasmid backbone and 40 fmol of each gBlock and/or PCR insert (as necessary). In a thermocycler, Golden Gate reactions were cycled 25 times: 90 s at 37 °C (for BsaI and SapI) or 42 °C (for BsmBI) followed by 3 minutes at 16 °C. After the 25 cycles, reactions were incubated at 37 °C (for BsaI and SapI) or 55 °C (for BsmBI) for 5 minutes, 80 °C for 10 minutes, and then held at 4 °C.

Golden Gate reactions were used to directly transform freshly prepared electrocompetent *S. oneidensis* strains. To prepare electrocompetent *S. oneidensis*, bacterial stocks stored in 22.5% glycerol at −80 °C were streaked onto LB agar plates and grown aerobically overnight at 30 °C. Single colonies were used to inoculate 5 mL of LB medium and grown aerobically in a shaking incubator (30 °C and 250 rpm). After 18 hours, the entire 5 mL culture was washed 3 times with sterile 10% glycerol at room temperature and concentrated to ∼300 *µ*L. 2 *µ*L of Golden Gate reaction was mixed with 30 *µ*L of concentrated electrocompetent *S. oneidensis*, transferred to a 1 mm electroporation cuvette, and electroporated at 1250 V. To recover electroporated cells, 250 *µ*L of LB was immediately added post-electroporation and cells were incubated/shaken at 30 °C and 250 rpm. After 2 h of recovery, 100 *µ*L of cell suspension was plated onto LB agar plates containing 25 *µ*g mL-1 kanamycin sulfate and incubated overnight at 30 °C to obtain single colonies (generally 5-100 colonies observed for 1-3 part assemblies). Single colonies were used to inoculate LB liquid medium containing 25 *µ*g mL^−1^ kanamycin sulfate and incubated/shaken overnight at 30 °C and 250 rpm. These cultures were used to generate 22.5% glycerol stocks, which were stored at −80 °C, and harvest assembled plasmid for Sanger sequencing (DNA Sequencing Facilities, University of Texas at Austin).

### RBS Library Design

Ribosome binding site (RBS) sequences with variable translation strength were computationally designed for each EET gene (*cymA, mtrA*, and *mtrC*) using the RBS Library Calculator (v2.0) from Salis et al.^43,44^ The calculator was utilized in design/search mode with the DNA sequence of each gene, the 16S rRNA sequence of *Shewanella oneidensis* MR-1, and the constant upstream region of each unique ribozyme after self-cleavage serving as software inputs (see Supplementary Table S2). The target minimum translation initiation rate (0.1 a.u.), target maximum translation initiation rate (1,000,000 a.u.), and target library size (64 members) were chosen such that output RBS sequences covered a broad translational space. One RBS sequence for *mtrC* (pCD24) was not designed using the RBS Library Calculator, and was instead derived from the BioBrick collection, B0032. The translational strength of this sequence (ΔG_total_ = −5.34 kcal mol^−1^) was calculated using the RBS Calculator (v2.0) using the same software inputs used for library construction. Translation strengths for each sequence were reported as ΔG_total_, which is related to translation initiation rate (r) according to the exponential relationship r ∝ exp(-βΔG_total_).

### Measurement of Fe(III) Reduction Kinetics by *S. oneidensis*

Except where noted, all Fe(III) reduction experiments were monitored *in situ* using 96-well plates that contained *S. oneidensis*, lactate, Fe(III) citrate, and ferrozine.^27^ The general protocol for an Fe(III) reduction setup was as follows. Bacterial stocks stored in 22.5% glycerol at −80 °C were streaked onto LB agar plates (for wild-type and knockout strains) or LB agar with 25 *µ*g mL^−1^ kanamycin (for plasmid-harboring strains). For anaerobic pregrowths, plates were brought into an anaerobic chamber (3% H_2_, balance N_2_, Coy) and single colonies were used to inoculate a pregrowth medium consisting of SBM supplemented with 100 mM HEPES, 1X trace mineral supplement (ATCC), 0.05% casamino acids, 20 mM sodium lactate as the electron donor, and 40 mM sodium fumarate as the electron acceptor. For dynamic expression and growth of strains absent of plasmid, this culture medium was inoculated and statically incubated in an anaerobic chamber at 30 °C for 18-24 hours. For steady-state expression experiments, this culture medium was inoculated and statically incubated in an anaerobic chamber at 30 °C for 6 hours. Afterwards, cultures were diluted 25-fold (10 *µ*L into 250 *µ*L final volume) into 96-well plates containing preinduction medium. Preinduction medium was identical to the SBM/lactate/fumarate pregrowth medium, but also contained varying concentrations of IPTG. These cultures were statically incubated in an anaerobic chamber at 30 °C for an additional 18-24 hours. Aerobic pregrowths were identical to the above except that single colonies were used to inoculate pregrowth medium that lacked 40 mM sodium fumarate and grown aerobically in a shaking incubator (250 rpm, 30 °C). All cultures were performed in biological triplicate where each culture was derived from a single picked colony. 25 *µ*g mL^−1^ kanamycin sulfate was added to pregrowth and preinduction media for strains harboring plasmids. Optical density for each overnight growth was monitored by pipetting 100 *µ*L (for preinduced samples) or 250 *µ*L cell suspension aliquots into 96-well plates and measuring absorbance at 600 nm using a BMG LABTECH CLARIOstar plate reader. The plate reader software converted measured optical density values into standard 10 mm cuvette readings based on the plate dimensions (96-well flat bottom, sterile, Greiner) and utilized liquid volume.

*In situ* Fe(III) reduction was set up within an anaerobic chamber using degassed media and solutions. Immediately prior to each experiment, 24 mg of ferrozine was dissolved in 12 mL of 2x SBM solution that was supplemented with 200 mM HEPES, 0.1% casamino acids, and 2X trace mineral supplement (ATCC). A master mix was generated that combined the 2X SBM/ferrozine with sodium lactate and kanamycin sulfate, such that the final concentrations of these components would be 1X casamino acid/trace mineral solution supplemented SBM, 1 mg mL^−1^ ferrozine, 20 mM sodium lactate, and 25 *µ*g mL^− 1^ kanamycin sulfate, within a final 250 *µ*L solution. 220 *µ*L aliquots of this master mix were pipetted into each 96-well plate well and subsequently mixed with 10 *µ*L of appropriate IPTG stock. Next, (un)induced *S. oneidensis* pregrowth was diluted 4-fold (25 *µ*L into 75 *µ*L SBM supplemented with kanamycin) in a separate 96-well plate. 10 *µ*L of this 4-fold dilution was used to inoculate the 96-well plate containing the Fe(III) reduction mixture (final dilution equal to 100-fold from overnight pregrowth). Lastly, 10 *µ*L of 125 mM Fe(III) citrate was added to each well such that the final concentration was 5 mM Fe(III). To generate an Fe(II) calibration curve, freshly prepared Fe(II) sulfate stocks dissolved in sterile water were also added to each 96-well plate. 10 *µ*L of each Fe(II) stock,10 *µ*L of sterile water, and 10 *µ*L abiotic SBM were mixed with 220 *µ*L of master mix such that the final concentration of Fe(II) standards ranged from 0 to 96 *µ*M Fe(II).

After addition of all plate components, the 96-well plate was sealed with a sterile and optically transparent sealing film (PCR-SP-S, AxySeal Scientific) and covered with a polystyrene plate lid (Eppendorf) that had silicone grease lining the edges. This plate was removed from the anaerobic chamber and placed within a BMG LABTECH CLARIOstar plate reader with temperature control set to 30 °C. Without shaking, the absorbance at 562 nm was measured every 10 minutes for at least 14 hours. For the ferrozine-free experiment, the Fe(III) reduction setup was identical to the above except that ferrozine was not dissolved in the 2X SBM. Instead, the 96-well plate was kept within an anaerobic chamber at 30 °C for the entirety of the experiment. Aliquots were taken and either acidified in equal volume 1 N HCl for measurement of Fe(II) levels or used for serial dilutions to perform colony counting on LB agar plates containing 25 *µ*g mL^−1^ kanamycin sulfate. Plated dilutions were grown aerobically overnight at 30 °C.

### Reduction Kinetics and Rate Constant Response Function Fitting

Derivations of the exponential and polynomial Fe(III) reduction models can be found in Supplementary Notes S1 and S2. Prior to kinetic fitting, the initial Fe(II) concentrations were subtracted from every timepoint within each replicate (i.e., 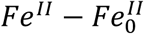). These subtracted Fe(II) concentrations, 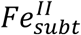, were then fit to either 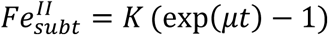 (exponential model) or 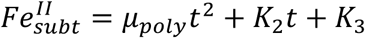 (polynomial model) to obtain *μ* and *μ*_*poly*_ rate constants, respectively. Response functions for Buffer gate circuits (fitted rate constants vs. IPTG) were modeled using a four-parameter activating Hill function 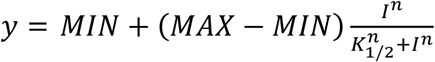. Response functions for NOT gate circuits were modeled using a four-parameter deactivating Hill function 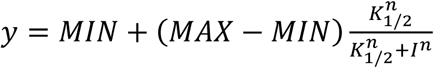. Individual replicate kinetic fitting was performed using least-squares regression with no weighting. Response function fitting was performed using least-squares regression weighting by 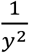 and each replicate *y* value was treated as an individual point. All fitting was performed in Prism 8.3 (GraphPad Software). Dynamic range was calculated as the ratio of induced (1500 *µ*M IPTG) and uninduced (0 *µ*M IPTG) fitted rate constants for each replicate, and the mean ± s.e.m. was determined.

