## Supplementary Information for "Tuning Extracellular Electron Transfer by *Shewanella oneidensis* Using Transcriptional Logic Gates"

### Note S1. Derivation of Monod-type Model for Fe(III) Reduction

*General relationship between cell growth and Fe(III) Reduction*

Monod-type models can be used to relate extracellular electron transfer (EET) and growth of electroactive microorganisms.[1] Generally, these models are of the form:

$$\frac{dN}{dt} = \mu N \quad (1)$$

$$\mu = \mu_{max} \frac{S}{K_S + S} \quad (2)$$

where $N$ is the concentration of cells in suspension, $\mu$ is the specific growth rate, $\mu_{max}$ is the maximum specific growth rate, and $S$ is the concentration of the growth rate-limiting substrate. Although in many situations the growth rate-limiting substrate is a carbon source,[2] it can also be an electron acceptor, which in our system is soluble $Fe^{III}$. In Monod-type models, $\mu$ is function of medium composition/cell-intrinsic physiology and, as discussed further below, can be assumed to remain constant.

Since cells utilize $Fe^{III}$ to generate further biomass, the following relationship arises in our system:

$$\frac{dN}{dt} = -Y \frac{dFe^{III}}{dt} \quad (3)$$

where $Y$ is the yield coefficient that relates substrate utilization to biomass generation. Combining (3) with (1) generates

$$\frac{dFe^{III}}{dt} = -\frac{\mu}{Y} N \quad (4)$$

Assuming constant $Y$ during metal reduction, (3) can also be integrated to

$$N = N_0 + Y(Fe_0^{III} - Fe^{III}) \quad (5)$$

By inserting (5) into (4), the following relationship is obtained

$$\frac{dFe^{III}}{dt} = -\frac{\mu}{Y}(N_0 + Y(Fe_0^{III} - Fe^{III})) \quad (6)$$

Equation (6) can be used to directly model microbial $Fe^{III}$ reduction kinetics and obtained fitted values for $Y$ and $\mu$. However, since the microbial reduction product $Fe^{II}$ is typically what is experimentally quantified, (6) can be rewritten using the following mass conservation relations

$$\frac{dFe^{III}}{dt} = -\frac{dFe^{II}}{dt} \quad (7)$$

$$Fe^{II} = Fe_0^{II} + Fe_0^{III} - Fe^{III} \quad (8)$$

to generate

$$\frac{dFe^{II}}{dt} = \frac{\mu}{Y}(N_0 + Y(Fe^{II} - Fe_0^{II})) \quad (9)$$

This differential equation can be integrated to

$$Fe^{II} - Fe_0^{II} = \frac{N_0}{Y}(\exp(\mu t) - 1) \quad (10)$$

Under steady-state gene expression, (10) is the exponential model that was used to fit kinetic data from our experiments and obtain values for $\mu$. We note that despite a generally strong fit of (10) to much of our experimental data, fitted values for $\mu$ appear higher than what has been experimentally measured for *Shewanella oneidensis* growth rates. Obtaining unrealistic growth rate values is a known issue when performing kinetic experiments with high $S_0/N_0$ ratios.[3] Since *S. oneidensis* also performs EET for energy maintenance in addition to cell growth, our fitted $\mu$ values are likely a combination of these factors. Thus, we elected to refer to $\mu$ as the fitted rate constant and not fitted growth rate.

*Comment on assumption of constant $\mu$*

In many conditions, $\mu$ can be assumed to remain constant when $S >> K_S$. Alternatively, $\mu$ can be assumed to remain constant when $S_0 >> \Delta S$, where $\Delta S$ is the change in $S$ over the course of the experiment. Our setup is the latter scenario, as we are only fitting $Fe^{II}$ kinetics when $Fe^{II}$ concentration ranges from 0 to 96 $\mu$M. As shown in Figure S6, we experimentally fit $K_S$ and determined its value to be 1889 $\mu$M, which compares favorably to previous measurements. (2) appears to accurately relate fitted rate constants to starting Fe(III) citrate concentration, which further validates that our kinetics follow a Monod-type relationship. Moreover, given the fitted $K_S$ value, our typical starting Fe(III) citrate concentration $S_0$ of ca. 5000 $\mu$M, and $\Delta S$ being at most 96 $\mu$M, $\mu$ deviates based on changing $S$ values by at most ca. 0.5%. This supports the validity of our assumption that fitted $\mu$ values are not affected by changing $Fe^{III}$ concentration under our experimental conditions.

*Relationship between fitted rate constant and EET gene expression*

The fitted rate constant from the Monod-type model controls the rate of microbial Fe(III) reduction within our system. When the cellular concentration of a single protein $P_{EET}$ rate-limits reduction of Fe(III), an additional relationship can be assumed for the Monod-type model that is similar to Michaelis-Menten enzyme kinetics.

$$\mu = \mu_{max} P_{EET} \frac{S}{K_S + S} \quad (11)$$

Since we expressed this EET protein in appropriate knockout strains of *S. oneidensis,* inducible control of $P_{EET}$ can control Fe(III) reduction. The dynamics of inducible protein expression are well studied and can be modeled using the following equations

$$\frac{dm_{EET}}{dt} = k_{tx}(MIN + (MAX - MIN)\frac{I^n}{K_{1/2}^n + I^n}) - d_m m_{EET} \quad (12)$$

$$\frac{dP_{EET}}{dt} = k_{tl} m_{EET} - d_P P_{EET} \quad (13)$$

where $m_{EET}$ is the cellular mRNA concentration of the target EET gene, $k_{tx}$ is a rate constant for this gene transcription, and $d_m$ is the degradation rate constant for this transcript. The expression $(MIN + (MAX - MIN)\frac{I^n}{K^n + I^n})$ is a quasi-empirical Hill function that describes LacI/IPTG regulation of EET gene transcription. The amount of EET gene mRNA generated depends on IPTG supplemented ($I$) and values for $MIN$, $MAX$, $n$, and $K_{1/2}$, which are typically obtained by model fitting. $k_{tl}$ is the rate constant for mRNA translation and $d_P$ is the protein degradation rate constant.

The steady-state approximation is frequently applied to (12), which causes $m_{EET}$ to follow a Hill function relationship

.

$$m_{EET,ss} = \frac{k_{tx}}{d_m}(MIN + (MAX - MIN)\frac{I^n}{K_{1/2}^n + I^n}) \quad (14)$$

After applying the steady-state approximation to (13), a linear relationship is derived between $m_{EET}$ and $P_{EET}$

$$P_{EET,ss} = \frac{k_{tl}}{d_P} m_{EET} \quad (15)$$

which can further be used with (14) to model $P_{EET,ss}$ as a Hill function in response to IPTG

$$P_{EET,ss} = \frac{k_{tl}}{d_P}\frac{k_{tx}}{d_m}(MIN + (MAX - MIN)\frac{I^n}{K_{1/2}^n + I^n}) \quad (16)$$

Given our assumption shown in (11), (16) predicts that $\mu$ can be described by a Hill function in response to IPTG. We note that $d_P$ is typically assumed to be equal to cellular growth rate. While the fitted $\mu$ values are related to growth rate, directly including them in $d_P$ likely leads to physiologically-unmeaningful dilution rates, since $\mu$ is a combination of growth rate and energetic maintenance requirements.[3] Based on comparable cell growth observed between induced/uninduced strains as shown in Figure S4, we primarily attribute drastic differences in fitted $\mu$ values (Figure 2) as a result of maintenance requirements.

**Note S2. Derivation of Polynomial Model for Fe(III) Reduction**

*Dynamic protein expression model*

The Monod-type model assumes that the cellular concentration of the Fe(III) reduction rate-limiting protein $P_{EET}$ is unchanging with time. This assumption is primarily valid during steady-state induction experiments. In contrast, $P_{EET}$ is expected to vary with time when performing dynamic expression experiments. This motivates the development of another model that captures the dynamics of protein concentration and can relate a fitted Fe(III) reduction parameter to $P_{EET}$.

While the Monod-type model assumes constant $P_{EET}$ and changing cell concentration, another limit may be examined where $P_{EET}$ varies with time and cell concentration remains relatively constant. This assumption affects (13), as the value of $d_P$, or essentially the cell growth rate, tends to 0:

$$\frac{dP_{EET}}{dt} = k_{tl} m_{EET,ss} \quad (17)$$

(17) implies that there is negligible growth-based dilution/protein degradation relative to the timescale of mRNA transcription/translation. (17) can be integrated to

$$P_{EET} = k_{tl} m_{EET,ss} t \quad (18)$$

which shows that $P_{EET}$ varies linearly with time and the steady-state concentration of EET gene mRNA $m_{EET}$. (18) can be further related to IPTG induction by replacing $m_{EET,ss}$ with (14)

$$P_{EET} = k_{tl} \frac{k_{tx}}{d_m} (MIN + (MAX - MIN) \frac{I^n}{K_{1/2}^n + I^n}) t \quad (19)$$

Thus, $P_{EET}$ is shown to vary linearly with time and the rate of $P_{EET}$ increase depends on supplemented IPTG concentrations.

*Connection of dynamic protein expression with Fe(III) reduction*

To relate Fe(III) reduction rate to $P_{EET}$, a variation of (4) can be assumed:

$$\frac{dFe^{III}}{dt} = -N_0(k_{EET}P_{EET} + k_{background}) \quad (20)$$

From (20), it is shown that $\frac{dFe^{III}}{dt}$ is proportional to the initial cell number and the cellular concentration of the rate-limiting EET protein $P_{EET}$. $k_{EET}$ is a rate constant that relates Fe(III) reduction to $P_{EET}$. (20) also captures background Fe(III) reduction that is independent of $P_{EET}$ via the rate constant $k_{background}$. Background Fe(III) reduction is assumed to be unchanging with time and proportional to initial cell number. (19) can be inserted into (20) to account for how $P_{EET}$ changes with time/IPTG concentration:

$$\frac{dFe^{III}}{dt} = -N_0(k_{EET}k_{tl}\frac{k_{tx}}{d_m}(MIN + (MAX - MIN)\frac{I^n}{K_{1/2}^n+I^n})t + k_{background}) \quad (21)$$

Integrating (21) and replacing Fe(III) variables with the mass conservation Fe(II) relations yields the second-order polynomial relationship

$$Fe^{II} - Fe_0^{II} = N_0\,(k_{EET}k_{tl}\frac{k_{tx}}{2d_m}(MIN + (MAX - MIN)\frac{I^n}{K_{1/2}^n+I^n}t^2 + k_{background}t) \quad (22)$$

Under dynamic gene expression, (22) is the polynomial model that was used to fit kinetic data from our experiments. Note that (22) predicts fitted constants preceding the $t^2$ term (which we refer to as $\mu_{poly}$) should follow a Hill function relationship based on supplemented IPTG concentration. For clarity, $\mu_{poly}$ is defined as

$$\mu_{poly} = N_0\,k_{EET}k_{tl}\frac{k_{tx}}{2d_m}(MIN + (MAX - MIN)\frac{I^n}{K_{1/2}^n+I^n}) \quad (23)$$

Together, (22) and (23) provide another model and fitted rate constant that can be used to parameterize inducible EET activity. Although we chose to examine dynamic protein expression under the no growth assumption to simplify the derived model, we note that growth is likely occurring. Thus, it may be possible to model reduction kinetics using other equations that combine aspects of (10) and (22), potentially through summed weighting.

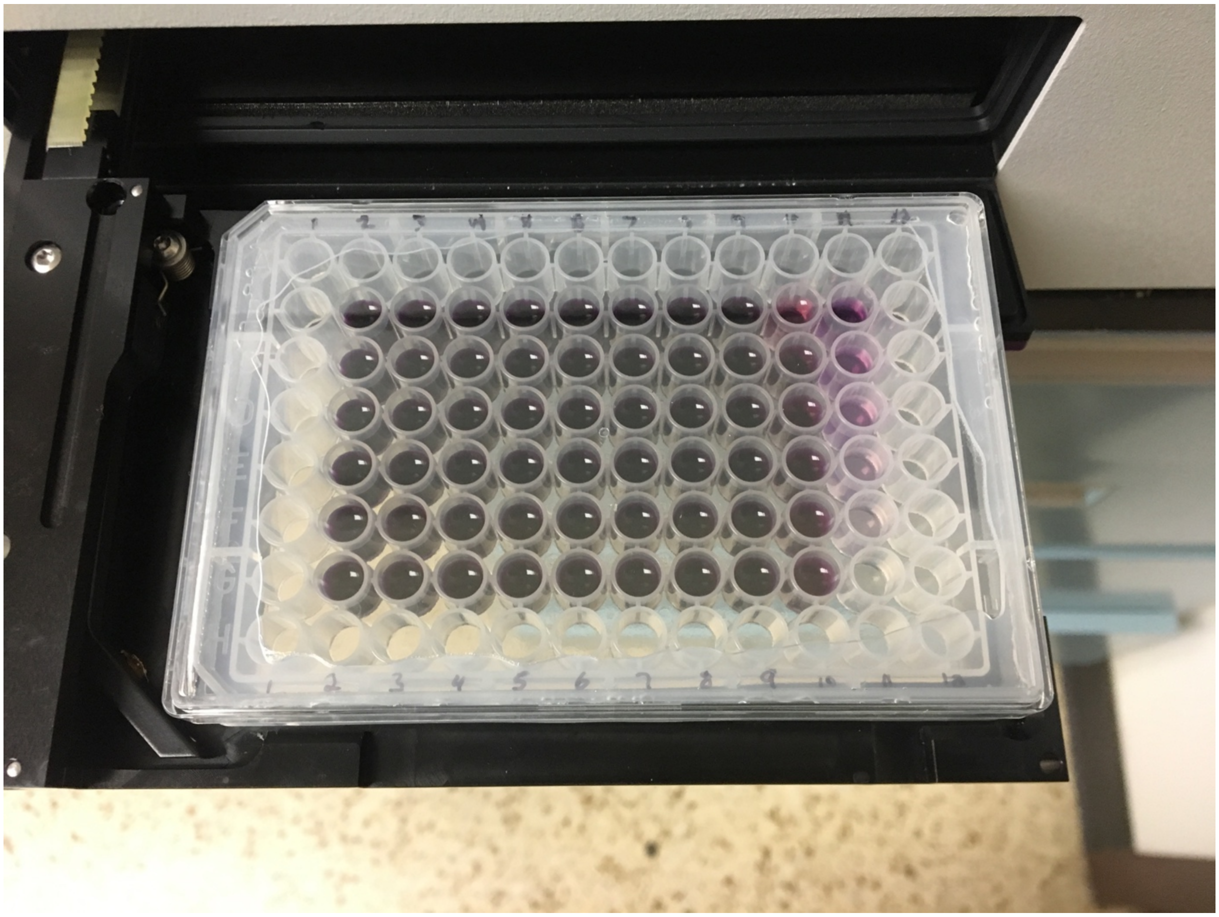

**Figure S1.** Anaerobically sealed 96-well plate within a plate reader after an 18-hour microbial Fe(III) reduction experiment. The dark purple color is a result of ferrozine in the growth medium complexing *S. oneidensis*-generated Fe(II). Labeled column 11 shows that the abiotic Fe(II) standards exhibit a range of purple hues after 18 hours, which suggests that the sealed plate maintains anaerobicity on the timescale of experiments.

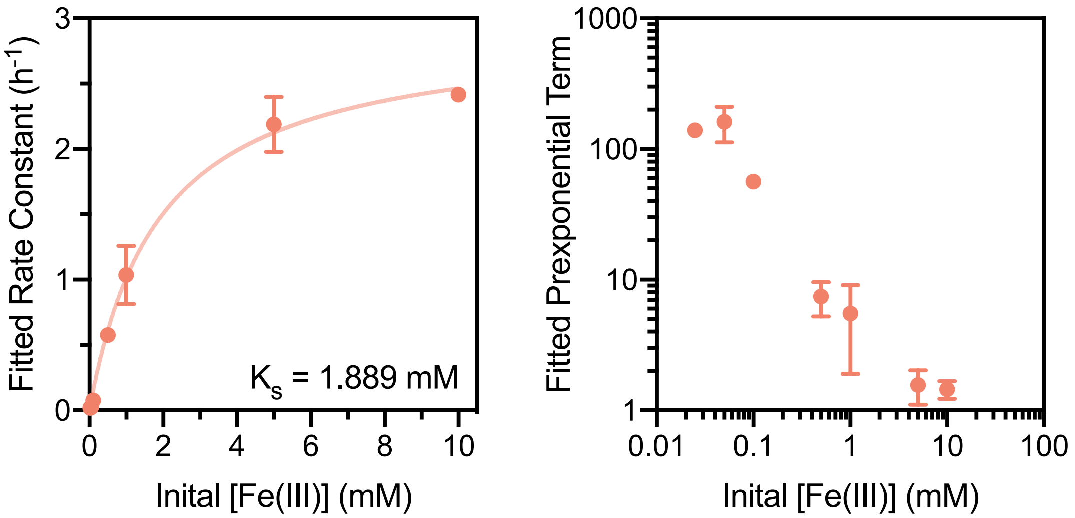


**Figure S2.** Inducible *mtrC* construct exhibits Monod-type behavior for fitted rate constant with varying initial Fe(III) concentration. A range of initial Fe(III) citrate concentrations were tested with the RBS-optimized *mtrC* construct (*S. oneidensis* Δ*mtrC*Δ*omcA*Δ*mtrF* +pCD24r1), which was induced with 1 mM IPTG. The solid line represents fitting of the exponential model rate constants from each tested Fe(III) concentration to equation (2), and $K_S$ was determined to be 1.889 mM (left). This value is comparable to previously measured values for *S. oneidensis* soluble Fe(III) Monod constants (0.028-6 mM)[4,5]. The prexponential term from equation (10) was also determined as a function of initial Fe(III) citrate concentration (right), and was generally invariant near the initial Fe(III) concentration used for most experiments (5 mM).

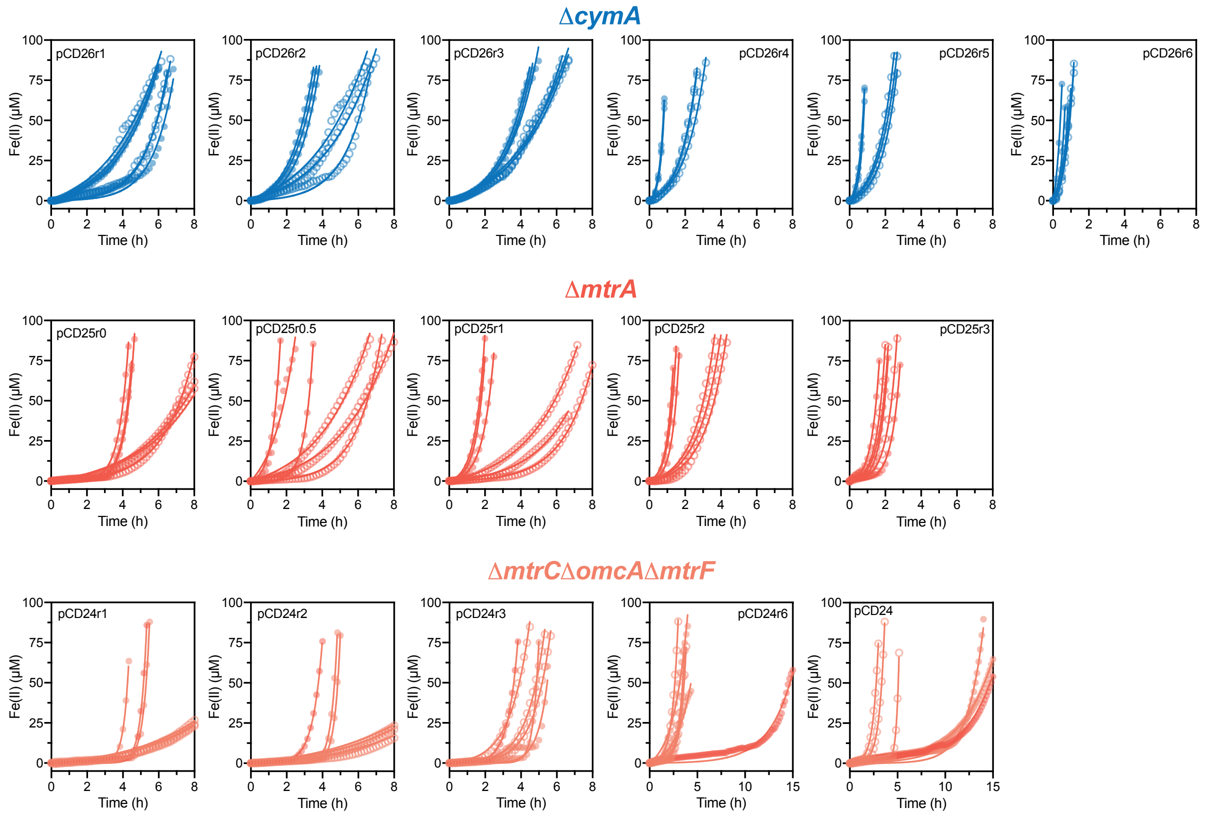


**Figure S3.** Raw kinetic data for RBS libraries. The fitted constants from this data are shown in main text Figure 3. Here, each graph shows induced (filled circles) and uninduced (open circles) replicates overlaid with respective exponential model fitted curves (solid lines). Induced samples were supplemented with 1500 $\mu$M IPTG and uninduced samples with 0 $\mu$M IPTG. Each row is a single RBS library (increasing translation strength from left to right), with the respective plasmid stated near the top of each graph. Above each row is the *S. oneidensis* knockout strain transformed by each RBS library member plasmid.

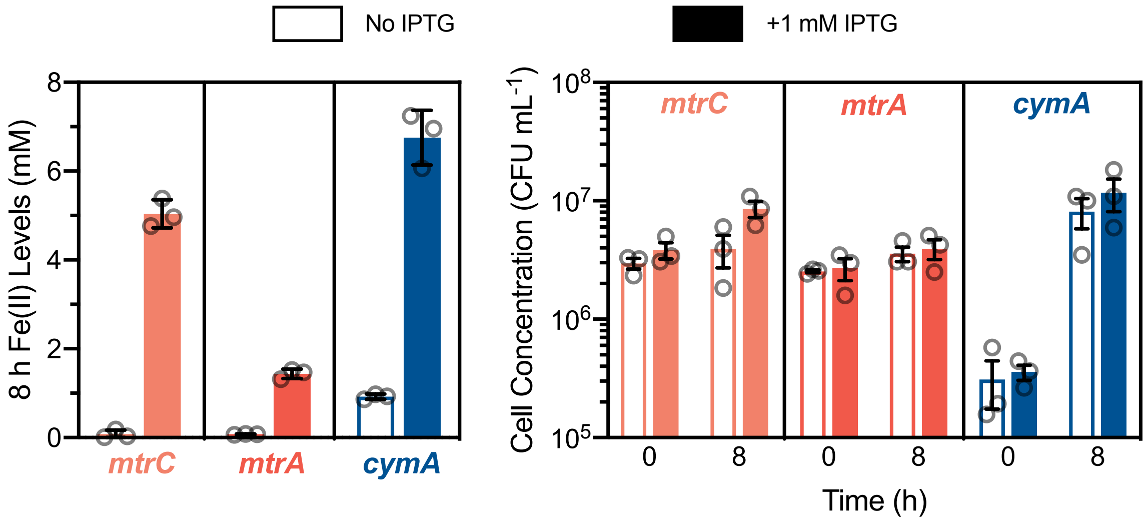


**Figure S4.** Concomitant Fe(III) reduction (left) and CFU counts (right) for RBS-optimized *mtrC* (*S. oneidensis* Δ*mtrC*Δ*omcA*Δ*mtrF*+pCD24r1), *mtrA* (*S. oneidensis* Δ*mtrA*+pCD25r0), and *cymA* (*S. oneidensis* Δ*cymA*+pCD26r4) constructs. The *mtrC* and *mtrA* constructs were pregrown anaerobically and the *cymA* construct was pregrown aerobically. Using the same Fe(III) reduction reaction conditions as in the main text experiments, 1 mL cultures were set up that omitted ferrozine in the Fe(III) reduction medium. Aliquots were acidified with equal volume 1 N HCl to quantify Fe(II) or serially diluted with SBM containing kanamycin for colony counting on LB agar plates. For Fe(III) reduction (left), data represent the mean ± s.d. for n = 3 biological replicates. For cell growth (right), data represent the mean ± s.e.m for n = 3 biological replicates.

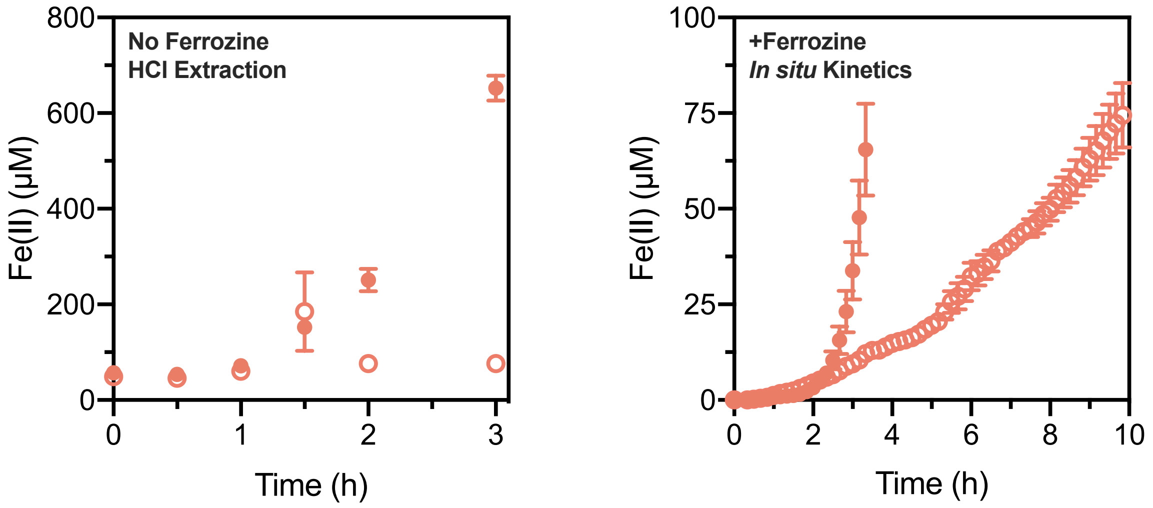


**Figure S5.** Early timescale Fe(III) reduction by the *mtrC* construct (*S. oneidensis* Δ*mtrC*Δ*omcA*Δ*mtrF*+pCD24r1). Experiments were performed either in the absence of ferrozine (left) or with ferrozine (right) in the growth medium. Other than the presence of ferrozine, Fe(III) reduction medium of both experiments was identical to typical reaction conditions detailed in the Materials and Methods (100-fold dilution from stationary-phase anaerobic pregrowth, 20 mM sodium lactate, 5 mM Fe(III) citrate).  For the left graph, experiments were performed within an anaerobic chamber at 30 °C, and aliquots were acidified with equal volume 1 N HCl for Fe(II) quantification. For the right graph, experiments were initially set up inside an anaerobic chamber and then moved to a BMG ClarioStar plate reader with temperature control set to 30 °C. Data for the right graph was obtained from Figure S3 and is included for comparison. Note that absolute Fe(II) levels are higher for HCl acidified samples, which likely is due to a combination of increased Fe(II) extraction[6] and quantification of intracellular Fe(II) from HCl-lysed cells. In both graphs, filled circles represent cultures supplemented with 1500 $\mu$M IPTG and open circles represent cultures with 0 $\mu$M IPTG. Data for both graphs represent the mean ± s.e.m. for n = 3 biological replicates.

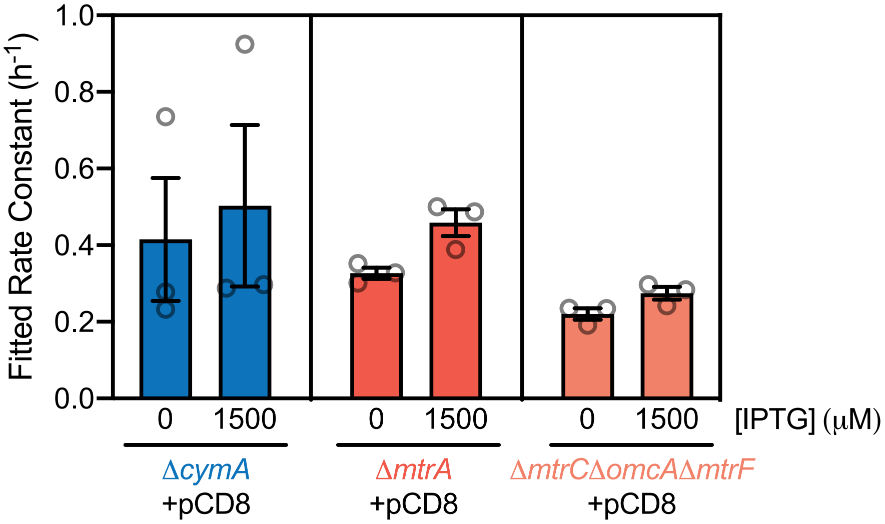


**Figure S6.** Fe(III) reduction by each Buffer gate strain background carrying the empty Buffer gate plasmid (pCD8). Each empty Buffer gate strain was tested in parallel with the respective Buffer gate shown in main text Figure 3. Induced samples were supplemented with 1500 $\mu$M IPTG and uninduced samples with 0 $\mu$M IPTG. Rate constants were determined using the exponential model. Data represent the mean ± s.e.m for n = 3 biological replicates.

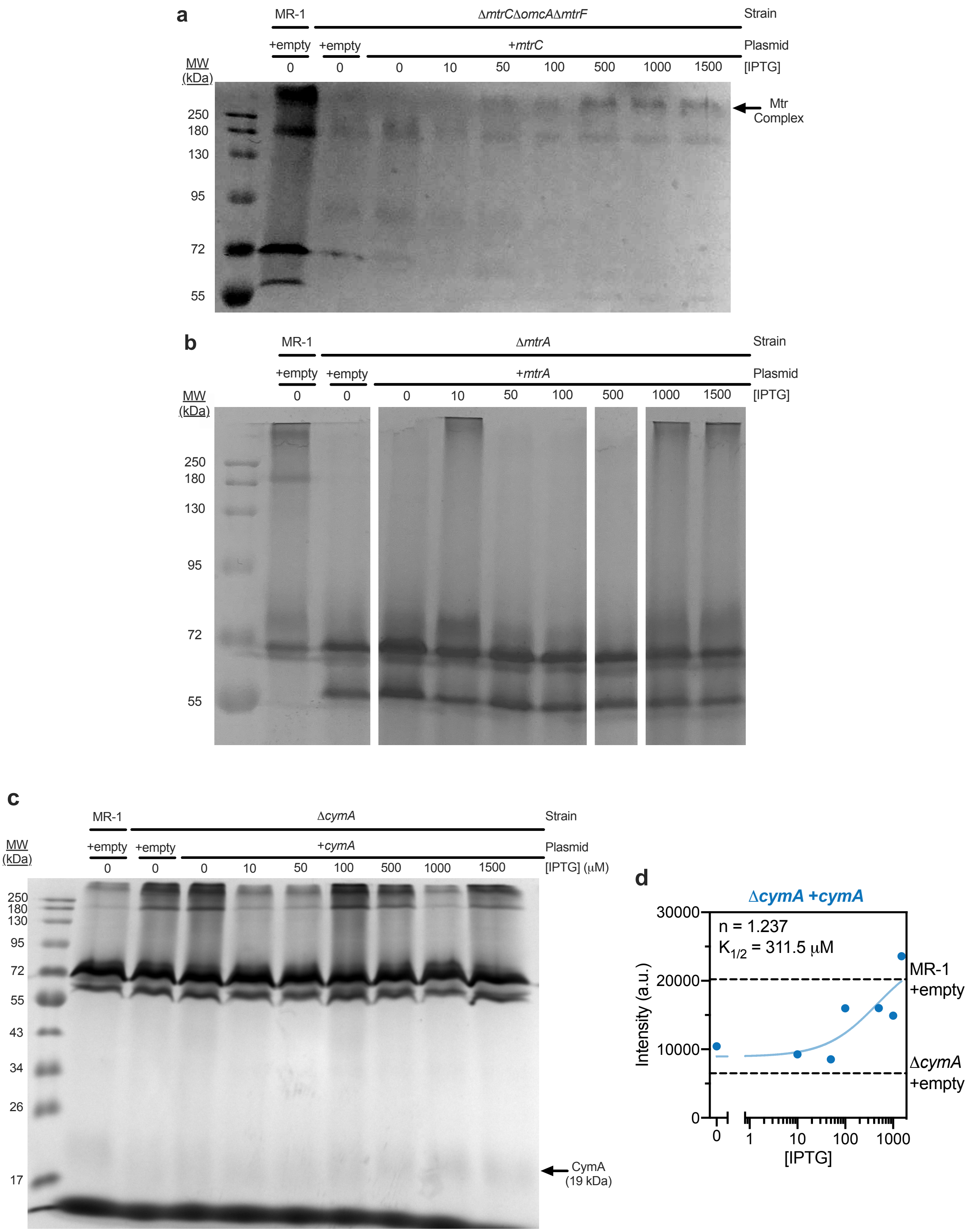


**Figure S7.** Heme staining of IPTG-induced EET gene expression constructs. (a) 10 *μ*g of protein was loaded into each lane for either the *mtrC* construct (*S. oneidensis* Δ*mtrC*Δ*omcA*Δ*mtrF* +pCD24r1) or positive (*S. oneidensis* MR-1 +pCD8) and negative

(*S. oneidensis* Δ*mtrC*Δ*omcA*Δ*mtrF*+pCD8) empty vector controls. (b) 25 μg of protein was loaded into each lane for either the *mtrA* construct (*S. oneidensis* Δ*mtrA*+pCD25r0) or positive (*S. oneidensis* MR-1+pCD8) and negative (*S. oneidensis* Δ*mtrA*+pCD8) empty vector controls. (c) 50 μg of protein was loaded into each lane for either the *cymA* construct (*S. oneidensis* Δ*cymA*+pCD26r4) or positive (*S. oneidensis* MR-1 +pCD8) and negative (*S. oneidensis* Δ*cymA*+pCD8) empty vector controls. (d) Densitometry of the CymA band in (c) denoted by the arrows was performed using the ImageJ gel analysis feature. The data was fit to an activating Hill function with the MIN and MAX values constrained by the intensities at 0 μM and 1500 μM IPTG, respectively. The general cell lysis/heme staining procedure was performed as follows. Briefly, 5 mL of cells that were induced for 18-24 hours with IPTG either aerobically (*cymA*) or anaerobically (*mtrC* and *mtrA*) were harvested by centrifugation at 3400 rcf for 30 minutes. The cells were washed once with 1 mL 1x Phosphate Buffered Saline (PBS) and the cell pellet was frozen at -80 °C. Prior to lysis, cells were thawed in 150 μL 1x PBS + 0.1% Triton, and the suspension was bath sonicated for 1 hour. Unlysed cell debris was spun down at 6000 rcf for 5 minutes, and the supernatant was separated, and protein content analyzed via Bradford assay. Heme staining was performed according to previous methods.

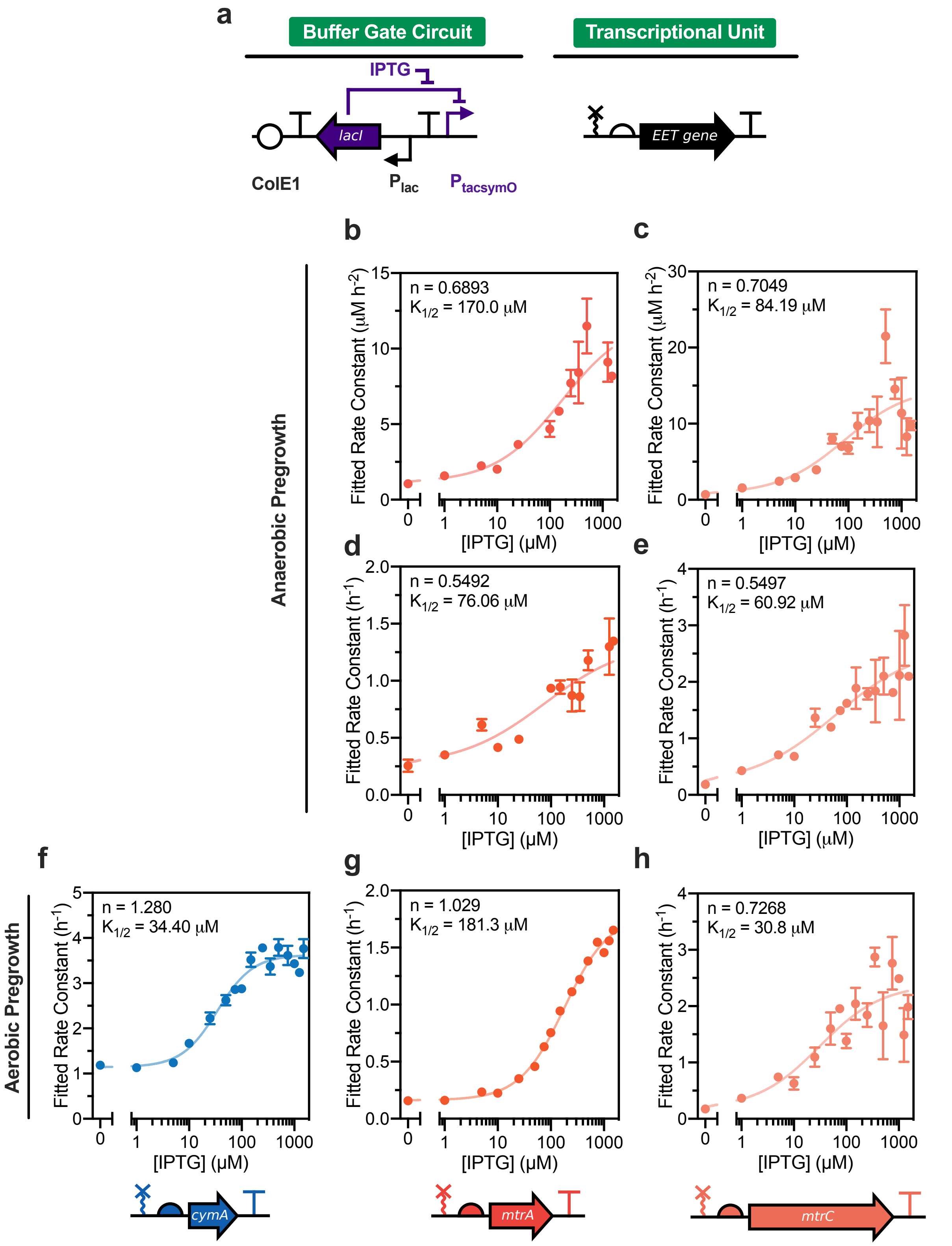


**Figure S8.** Buffer gate response functions under dynamic expression for *cymA*, *mtrA*, and *mtrC* constructs using Monod and polynomial parameters. (a) Diagram of Buffer gate circuit used to control each EET gene transcriptional output. (b) and (d) Fe(III) reduction response functions generated by anaerobically pregrown *S. oneidensis* Δ*mtrA* expressing the *mtrA* Buffer gate circuit (pCD25r0). Kinetics were fit using the (b) polynomial model or (d) exponential model. (c) and (e) Fe(III) reduction response function generated by anaerobically pregrown *S. oneidensis* Δ*mtrC*Δ*omcA*Δ*mtrF* expressing the *mtrC* Buffer gate circuit (pCD24r1). Kinetics were fit using the (c) polynomial model or (e) exponential model. Aerobic pregrowth Fe(III) reduction response function generated by the (f) *cymA*, (g) *mtrA*, and (h) *mtrC* constructs. Kinetics for (f)-(h) were fit using the exponential model. In these experiments, IPTG was supplemented immediately prior to measuring Fe(III) reduction kinetics. The solid line and Hill parameters shown in each graph were obtained by fitting individual replicates to a four-parameter activating Hill function with weighted error. To improve Hill function fitting for (d), the minimum and maximum Hill function values were constrained by the mean fitted rate constants at 0 and 1500 $\mu$M IPTG, respectively. All data represent the mean ± s.e.m. of fitted rate constants for n = 3 biological replicates.

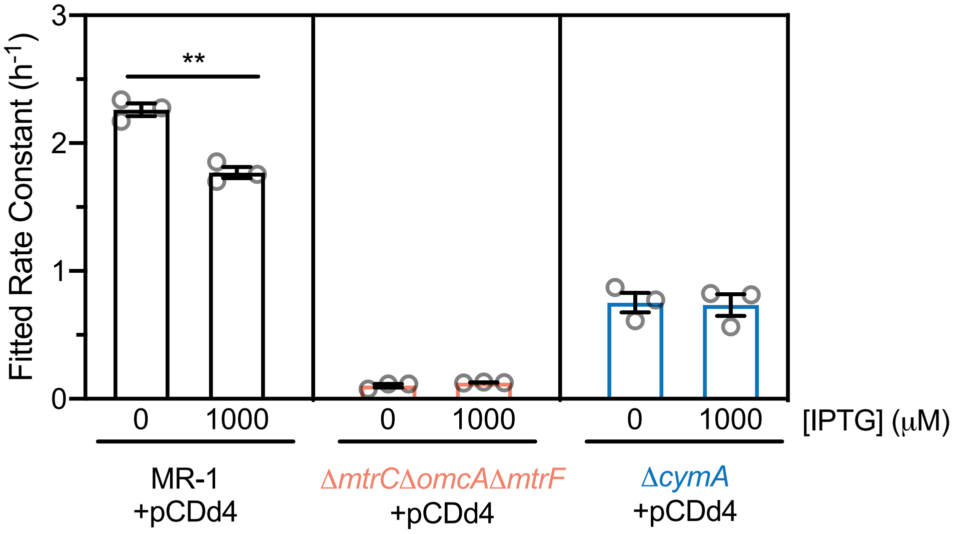


**Figure S9.** NOT gate empty vector controls with MR-1 and knockout strains. The NOT gate circuit lacking any transcriptional output gene (pCDd4) was used to transform *S. oneidensis* strains MR-1, Δ*mtrC*Δ*omcA*Δ*mtrF*, and Δ*cymA*. MR-1 and the Δ*mtrC*Δ*omcA*Δ*mtrF* strains were pregrown anaerobically. The Δ*cymA* strain was pregregrown aerobically. All strains were preinduced for 24 hours with IPTG prior to analyzing Fe(III) reduction kinetics. Kinetics were fit to the exponential model to obtain fitted rate constants. Data represent mean mean ± s.e.m. for n = 3 biological replicates. **p = 0.0018.

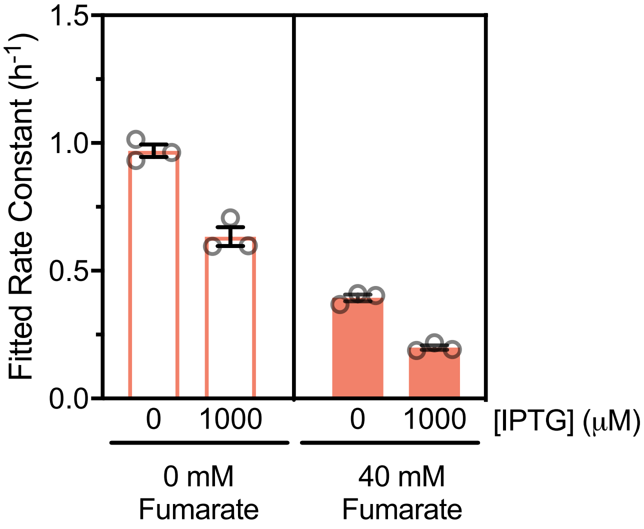


**Figure S10.** Dynamic NOT gate activity for *mtrC* construct with 0 and 40 mM sodium fumarate. *S. oneidensis* Δ*mtrC*Δ*omcA*Δ*mtrF* was pregrown anaerobically in the absence of IPTG. 0 or 1000 $\mu$M IPTG was added to microbially Fe(III) reduction reactions immediately prior to kinetic measurements. Additionally, experiments were supplemented with either 0 or 40 mM sodium fumarate during the Fe(III) reduction. Kinetics were fit to the exponential model to obtain fitted rate constants. Data represent mean ± s.e.m. for n = 3 biological replicates.

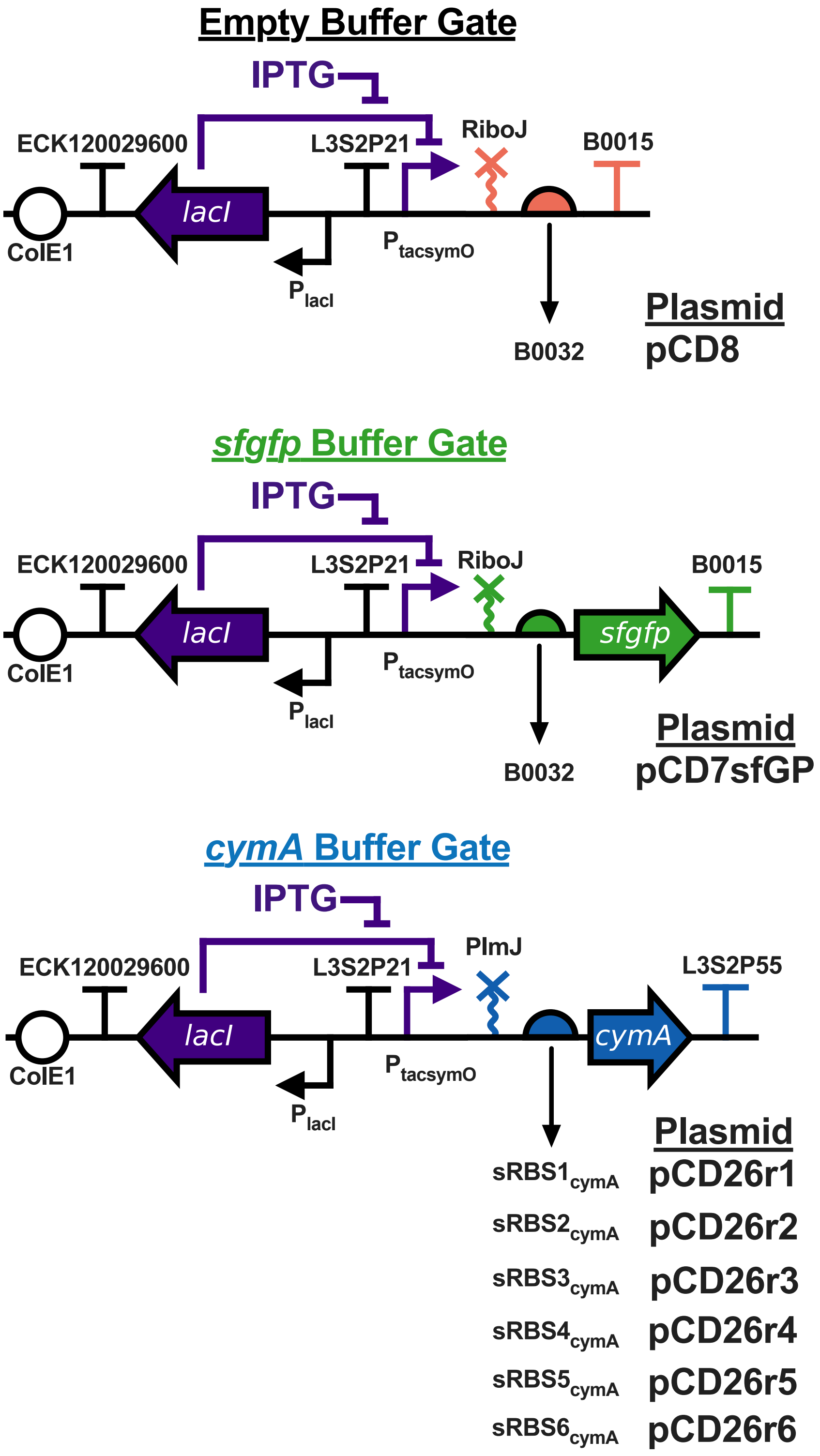
Empty Buffer Gate
IPTG
ECK120029600
L3S2P21
RiboJ
B0015
lacI
ColE1
P tacsymO
P lacI
B0032
Plasmid
pCD8
sfgfp Buffer Gate
IPTG
ECK120029600
L3S2P21
RiboJ
B0015
lacI
sfgfp
ColE1
P tacsymO
P lacI
B0032
Plasmid
pCD7sfGP
cymA Buffer Gate
IPTG
ECK120029600
L3S2P21
PlmJ
L3S2P55
lacI
cymA
ColE1
P tacsymO
P lacI
Plasmid
sRBS1 cymA pCD26r1
sRBS2 cymA pCD26r2
sRBS3 cymA pCD26r3
sRBS4 cymA pCD26r4
sRBS5 cymA pCD26r5
sRBS6 cymA pCD26r6

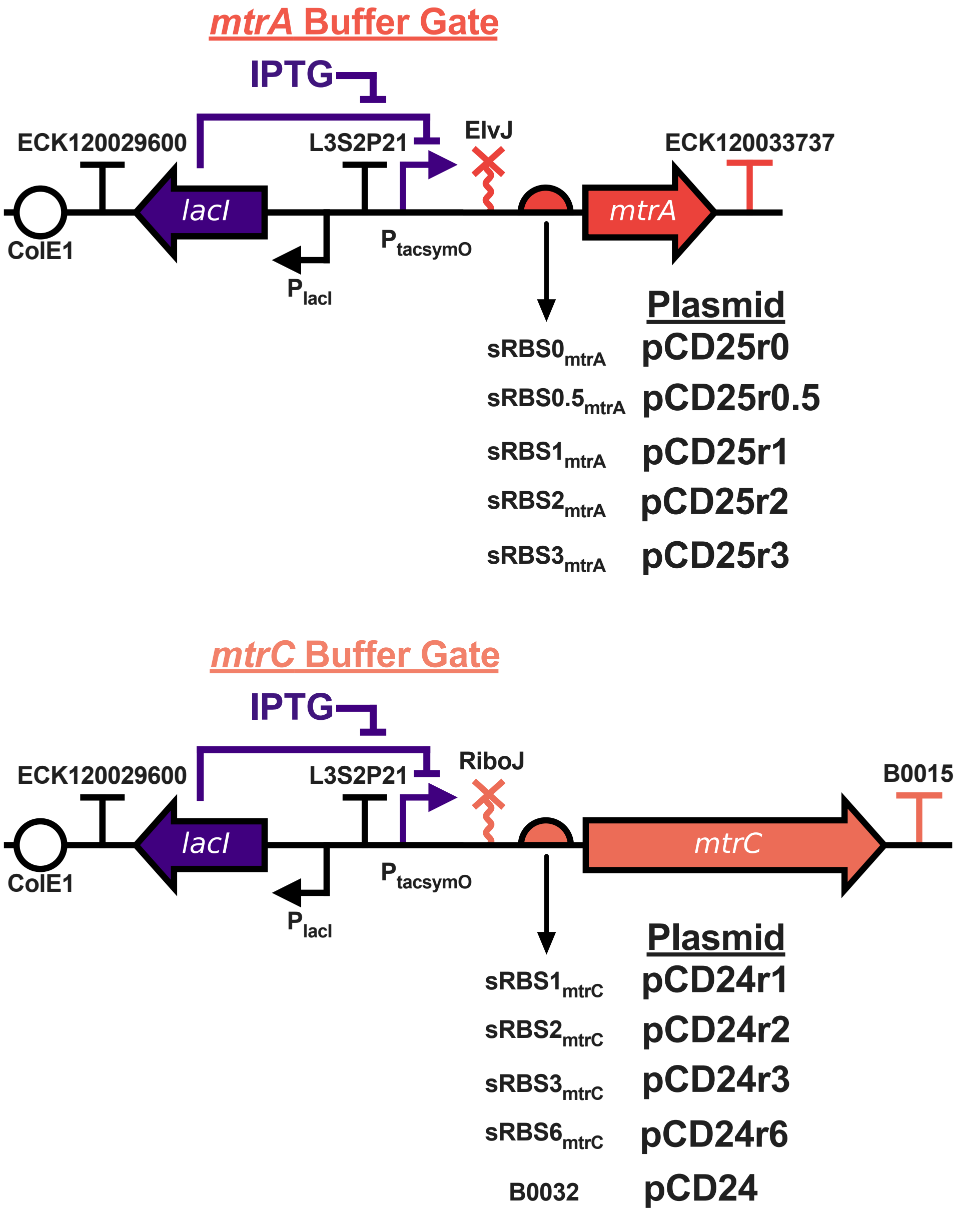


**Figure S10.** Gene circuit maps for the empty, *sfgfp*, *cymA*, *mtrA*, and *mtrC* Buffer gate plasmids. Outside of the flanking terminators, all vectors are identical and contain the ColE1 origin of replication and a kanamycin resistance marker.

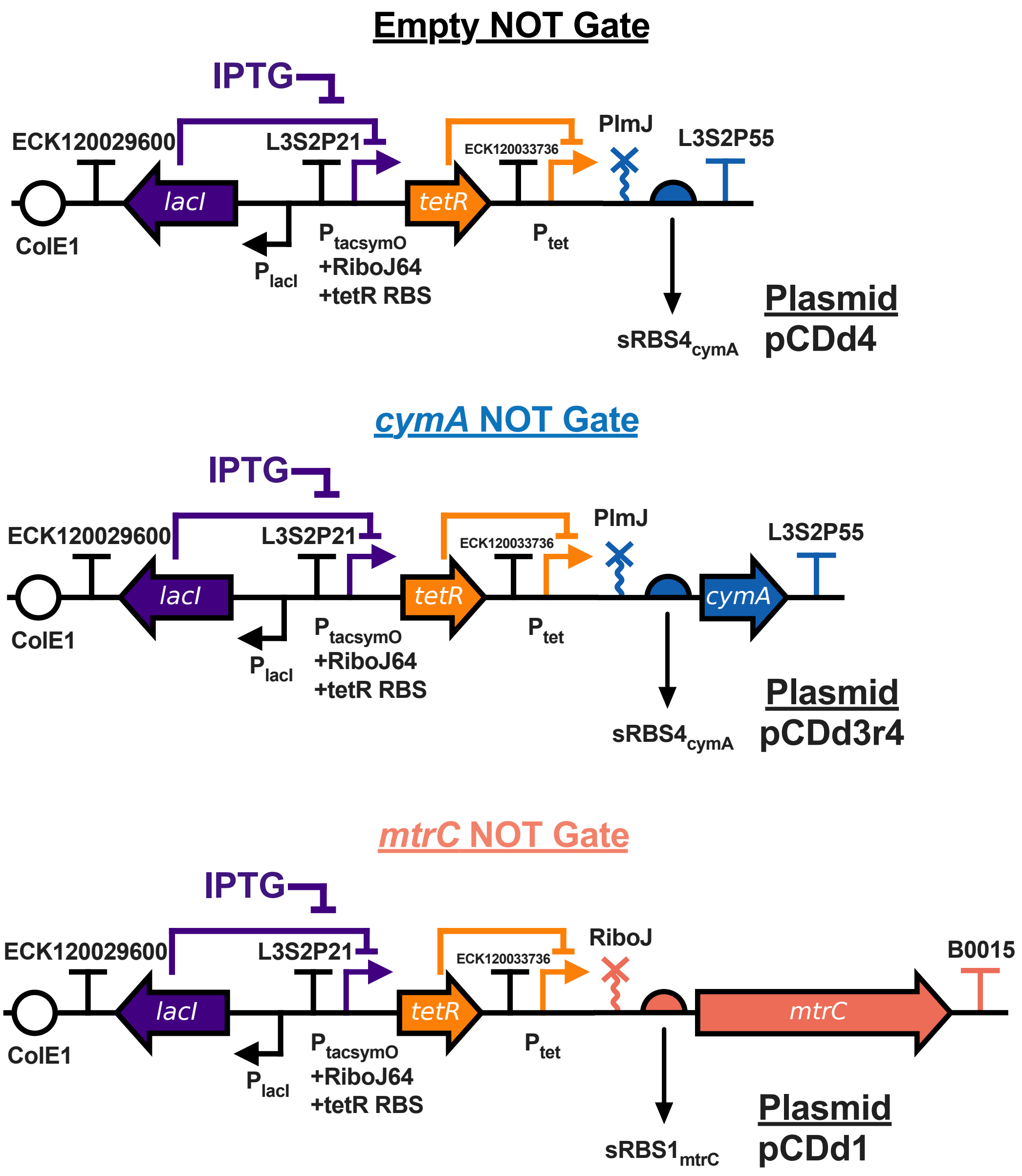


**Figure S11.** Gene circuit maps for the empty, *cymA*, and *mtrC* NOT gate plasmids. Outside of the flanking terminators, all vectors are identical and contain the ColE1 origin of replication and a kanamycin resistance marker.

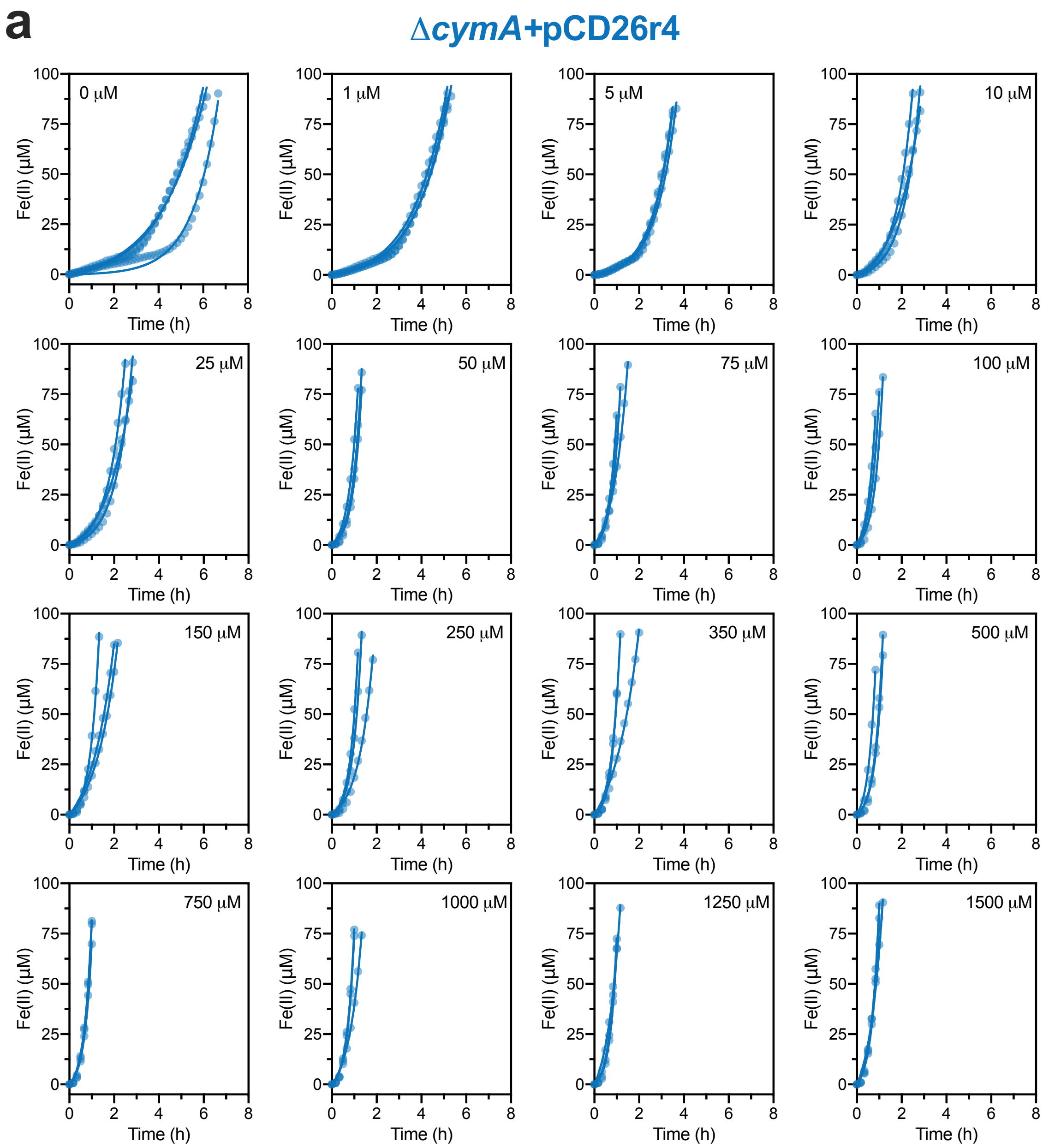
a
ΔcymA+pCD26r4
0 μM
1 μM
5 μM
10 μM
25 μM
50 μM
75 μM
100 μM
150 μM
250 μM
350 μM
500 μM
750 μM
1000 μM
1250 μM
1500 μM
Fe(II) (μM)
Time (h)

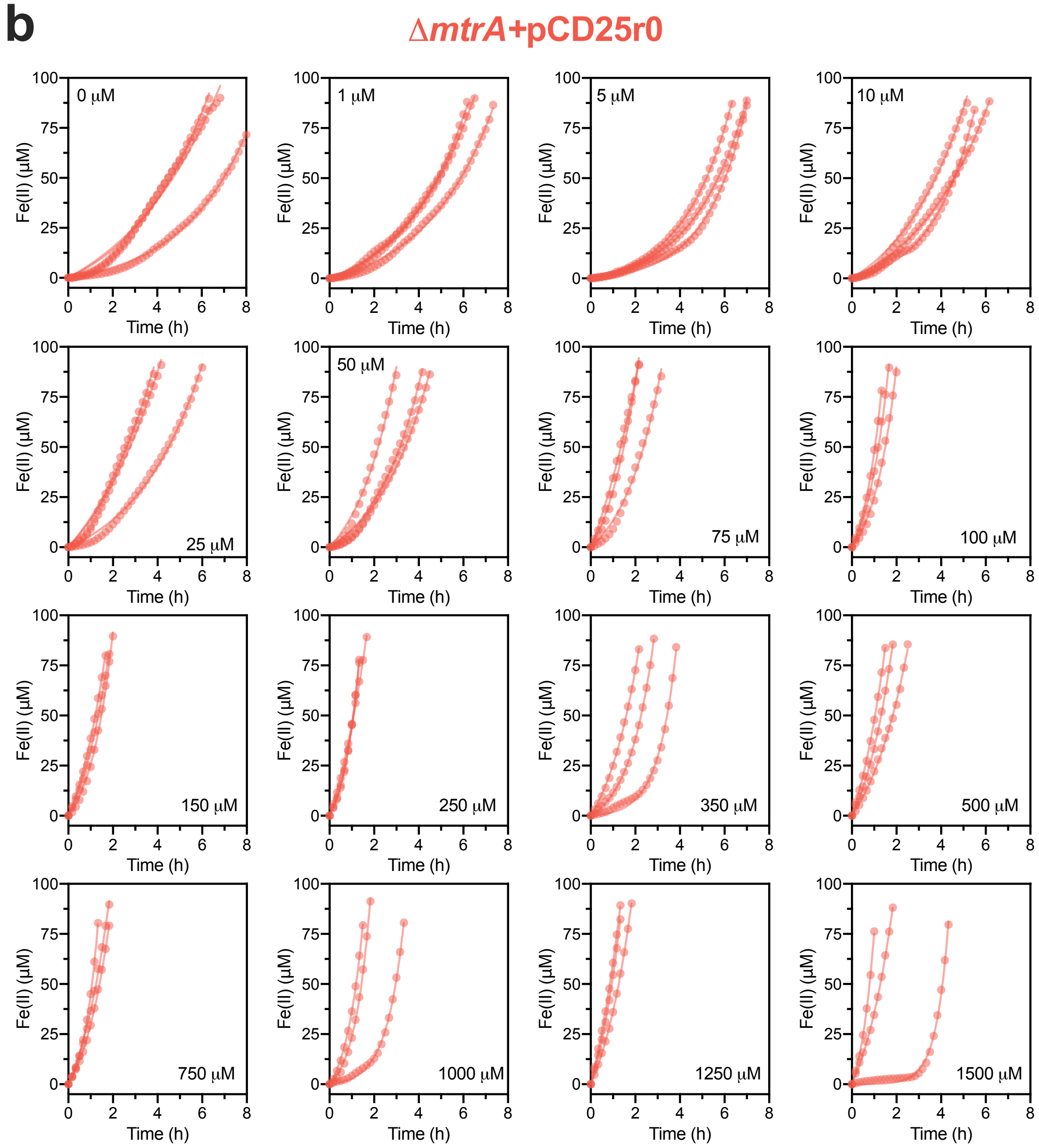
b
ΔmtrA+pCD25r0
Fe(II) (μM)
Time (h)
0 μM
1 μM
5 μM
10 μM
25 μM
50 μM
75 μM
100 μM
150 μM
250 μM
350 μM
500 μM
750 μM
1000 μM
1250 μM
1500 μM

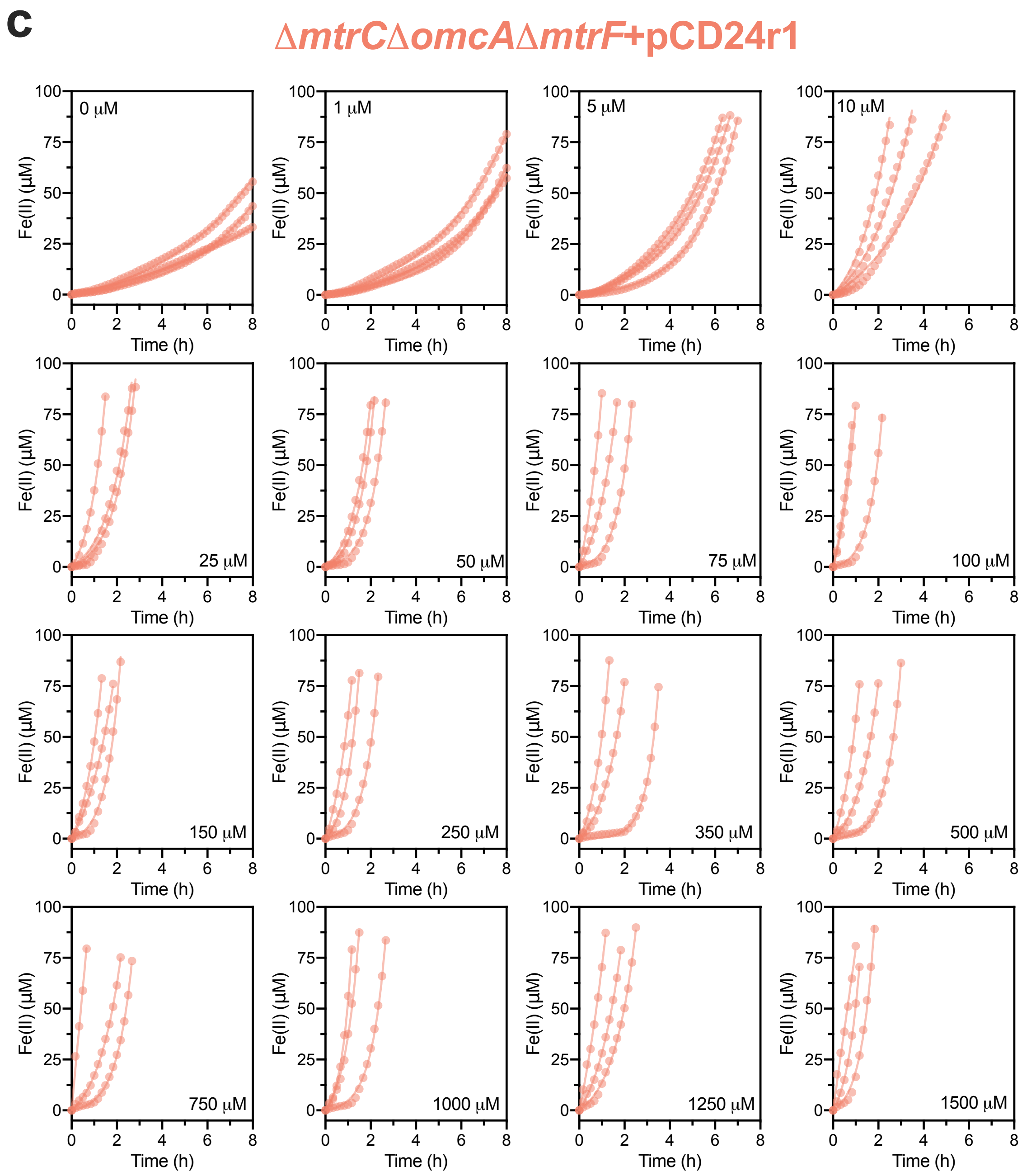


**Figure S12.** Raw kinetic data and replicates for main text Figure 4. (a) Raw kinetic data and overlaid fitted curves for Figure 4c. (b) Raw kinetic data and overlaid fitted curves for Figure 4d. (c) Raw kinetic data and overlaid fitted curves for Figure 4e. IPTG concentrations are stated in the corner of each graph.

**Table S1.** Bacterial strains and plasmids used in this study.

| Strain or plasmid | Description/Genotype | Reference or source |
|---|---|---|
| ***S. oneidensis* Strains (+plasmid)** | | |
| MR-1 | MR-1 (ATCC700550), wild-type strain | American-Type Culture Collection |
| JG596 | Lacks outer membrane cytochromes MtrC, OmcA, and MtrF; Δ*mtrC*Δ*omcA*Δ*mtrF* | [12] |
| Δ*cymA* | Lacks inner membrane cytochrome CymA | Jeffrey Gralnick, U. of Minnesota |
| Δ*mtrA* | Lacks periplasmic cytochrome, MtrA | This work |
| MR-1+pCD7sfGFP | Wild-type with *sfgfp* Buffer gate | This work |
| MR-1+pCD8 | Wild-type with empty Buffer gate | This work |
| MR-1+pCDd4 | Wild-type with empty NOT gate | This work |
| Δ*cymA*+pCD8 | Δ*cymA* with empty Buffer gate | This work |
| Δ*cymA*+pCD26r1 | Δ*cymA* with *cymA* Buffer gate ($sRBS1_{cymA}$) | This work |
| Δ*cymA*+pCD26r2 | Δ*cymA* with *cymA* Buffer gate ($sRBS2_{cymA}$) | This work |
| Δ*cymA*+pCD26r3 | Δ*cymA* with *cymA* Buffer gate ($sRBS3_{cymA}$) | This work |
| Δ*cymA*+pCD26r4 | Δ*cymA* with *cymA* Buffer gate ($sRBS4_{cymA}$) | This work |
| Δ*cymA*+pCD26r5 | Δ*cymA* with *cymA* Buffer gate ($sRBS5_{cymA}$) | This work |
| Δ*cymA*+pCD26r6 | Δ*cymA* with *cymA* Buffer gate ($sRBS6_{cymA}$) | This work |
| Δ*cymA*+pCDd4 | Δ*cymA* with empty NOT gate | This work |
| Δ*cymA*+pCDd3r4 | Δ*cymA* with *cymA* NOT gate ($sRBS4_{cymA}$) | This work |
| Δ*mtrA*+pCD8 | Δ*mtrA* with empty Buffer gate | This work |
| Δ*mtrA*+pCD25r0 | Δ*mtrA* with *mtrA* Buffer gate ($sRBS0_{mtrA}$) | This work |
| Δ*mtrA*+pCD25r0.5 | Δ*mtrA* with *mtrA* Buffer gate ($sRBS0.5_{mtrA}$) | This work |
| Δ*mtrA*+pCD25r1 | Δ*mtrA* with *mtrA* Buffer gate ($sRBS1_{mtrA}$) | This work |
| Δ*mtrA*+pCD25r2 | Δ*mtrA* with *mtrA* Buffer gate ($sRBS2_{mtrA}$) | This work |
| Δ*mtrA*+pCD25r3 | Δ*mtrA* with *mtrA* Buffer gate ($sRBS3_{mtrA}$) | This work |
| JG596+pCD8 | JG596 with empty Buffer gate | This work |

| | | |
|---|---|---|
| JG596+pCD24r1 | JG596 with *mtrC* Buffer gate (sRBS1$_{mtrC}$) | This work |
| JG596+pCD24r2 | JG596 with *mtrC* Buffer gate (sRBS2$_{mtrC}$) | This work |
| JG596+pCD24r3 | JG596 with *mtrC* Buffer gate (sRBS3$_{mtrC}$) | This work |
| JG596+pCD24r6 | JG596 with *mtrC* Buffer gate (sRBS6$_{mtrC}$) | This work |
| JG596+pCD24 | JG596 with *mtrC* Buffer gate (B0032) | This work |
| JG596+pCDd4 | JG596 with empty NOT gate | This work |
| JG596+pCDd1 | JG596 with *mtrC* NOT gate (sRBS1$_{mtrC}$) | This work |
| **Plasmids** | | |
| pCD7sfGFP | *sfgfp* Buffer gate | This work |
| pCD8 | Empty Buffer gate | This work |
| pCD24r1 | *mtrC* Buffer gate (sRBS1$_{mtrC}$) | This work |
| pCD24r2 | *mtrC* Buffer gate (sRBS2$_{mtrC}$) | This work |
| pCD24r3 | *mtrC* Buffer gate (sRBS3$_{mtrC}$) | This work |
| pCD24r6 | *mtrC* Buffer gate (sRBS6$_{mtrC}$) | This work |
| pCD24 | *mtrC* Buffer gate (B0032) | This work |
| pCD25r0 | *mtrA* Buffer gate (sRBS0$_{mtrA}$) | This work |
| pCD25r0.5 | *mtrA* Buffer gate (sRBS0.5$_{mtrA}$) | This work |
| pCD25r1 | *mtrA* Buffer gate (sRBS1$_{mtrA}$) | This work |
| pCD25r2 | *mtrA* Buffer gate (sRBS2$_{mtrA}$) | This work |
| pCD25r3 | *mtrA* Buffer gate (sRBS3$_{mtrA}$) | This work |
| pCD26r1 | *cymA* Buffer gate (sRBS1$_{cymA}$) | This work |
| pCD26r2 | *cymA* Buffer gate (sRBS2$_{cymA}$) | This work |
| pCD26r3 | *cymA* Buffer gate (sRBS3$_{cymA}$) | This work |
| pCD26r4 | *cymA* Buffer gate (sRBS4$_{cymA}$) | This work |
| pCD26r5 | *cymA* Buffer gate (sRBS5$_{cymA}$) | This work |
| pCD26r6 | *cymA* Buffer gate (sRBS6$_{cymA}$) | This work |
| pCDd4 | Empty NOT gate | This work |
| pCDd1 | *mtC* NOT gate (sRBS1$_{mtrC}$) | This work |
| pCDd3r4 | *cymA* NOT gate (sRBS4$_{cymA}$) | This work |

**Table S2.** Strain/Plasmid Information for RBS Library

| Strains | Plasmid | EET Gene RBS | Predicted RBS Strength (kcal mol$^{-1}$) |
|---|---|---|---|
| JG596 | pCD24r1 | sRBS1$_{mtrC}$ | 18.6 |
| JG596 | pCD24r2 | sRBS2$_{mtrC}$ | 16.0 |
| JG596 | pCD24r3 | sRBS3$_{mtrC}$ | 10.6 |
| JG596 | pCD24r6 | sRBS6$_{mtrC}$ | 3.55 |
| JG596 | pCD24 | B0032 | -5.34 |
| Δ*mtrA* | pCD25r0 | sRBS0$_{mtrA}$ | 16.4 |
| Δ*mtrA* | pCD25r0.5 | sRBS0.5$_{mtrA}$ | 13.4 |
| Δ*mtrA* | pCD25r1 | sRBS1$_{mtrA}$ | 9.15 |
| Δ*mtrA* | pCD25r2 | sRBS2$_{mtrA}$ | 4.00 |
| Δ*mtrA* | pCD25r3 | sRBS3$_{mtrA}$ | -1.11 |
| Δ*cymA* | pCD26r1 | sRBS1$_{cymA}$ | 23.1 |
| Δ*cymA* | pCD26r2 | sRBS2$_{cymA}$ | 18.2 |
| Δ*cymA* | pCD26r3 | sRBS3$_{cymA}$ | 13.3 |
| Δ*cymA* | pCD26r4 | sRBS4$_{cymA}$ | 8.19 |
| Δ*cymA* | pCD26r5 | sRBS5$_{cymA}$ | 3.06 |
| Δ*cymA* | pCD26r6 | sRBS6$_{cymA}$ | -1.89 |

**Table S3.** Genetic parts/sequences used to construct the plasmids in this study. Underline indicates added insulator sequence.

| Genetic Part | DNA Sequence (5' to 3') |
|---|---|
| **Promoters** | |
| $P_{tacsymO}$[7] | TGTTGACAATTAATCATCGGCTCGTATAATGTGTGGAATTGTGAGCGCTCACAATTCTATGGACTATGTTT |
| $P_{lacI}$ | GCGGCGCGCCATCGAATGGCGCAAAACCTTTCGCGGTATGGCATGATAGCGCCCGGAAGAGAGTCAATTCAGGGTGGTGAAT |
| $P_{tet}$[8] | TACTCCACCGTTGGCTTTTTTCCCTATCAGTGATAGAGATTGACATCCCTATCAGTGATAGAGATAATGAGCAC |
| **Ribosome Binding Sites** | |
| B0032 | TCACACAGGAAAGTACTAG |
| sRBS1$_{mtrC}$ | GGGGAAAAACAGCAGTGCGAT |
| sRBS2$_{mtrC}$ | GGAGAAAAACAGCAGTGCGAT |
| sRBS3$_{mtrC}$ | CGGCAAAAACAGCAGTACGAA |
| sRBS6$_{mtrC}$ | CGGGAAAAACAGGAGTGCGAA |
| sRBS0$_{mtrA}$ | ATAGGCGGCTTCATTGACGGTCCCA |
| sRBS0.5$_{mtrA}$ | ATAGGCGGCTTCACTGACGGTCCCA |
| sRBS1$_{mtrA}$ | TTACCCTAAGCTTCAGTAGGCGTTC |
| sRBS2$_{mtrA}$ | ATAGGCTGCTTCGCTGAGGGTCCCA |
| sRBS3$_{mtrA}$ | ATAGACTGCTTCGCTGAGGGTCCCA |
| sRBS1$_{cymA}$ | TTCGCTTTGGGTTTTTATGTAGCACGCA |
| sRBS2$_{cymA}$ | TTCGCTTTGGGTTTTTATGTAGGACGCA |
| sRBS3$_{cymA}$ | TTCCCTTTGGGTTTTTAAGTAGCACGCA |
| sRBS4$_{cymA}$ | TTCGCTTTGGGTTTTTAAGGAGGACGCA |
| sRBS5$_{cymA}$ | TTCCCTTTTGGTTTTTACGGAGCACGGA |
| sRBS6$_{cymA}$ | TTCCCTTTTGGTTTTTACGGAGGACGGA |
| tetR RBS[9] | CTATGGACTATGTTTTCACACAGGAAAGGCCTCG |
| **Terminators** | |
| ECK120029600[10] | TTCAGCCAAAAAACTTAAGACCGCCGGTCTTGTCCACTACCTTGCAGTAATGCGGTGGACAGGATCGGCGGTTTTCTTTTCTCTTCTCAA |
| L3S2P21[10] | TCGGTACCAAATTCCAGAAAAGAGGCCTCCCGAAAGGGGGGCCTTTTTTCGTTTTGGTCC |
| B0015 | TAATCTAGACCAGGCATCAAATAAAACGAAAGGCTCAGTCGAAAGACTGGGCCTTTCGTTTTATCTGTTGTTTGTCGGTGAACGCTCTCTACTAGAGTCACACTGGCTCACCTTCGGGTGGGCCTTTCTGCGTTTATA |
| ECK120033737[10] | GGAAACACAGAAAAAAGCCCGCACCTGACAGTGCGGGCTTTTTTTTTCGACCAAAGG |
| L3S2P55[10] | CTCGGTACCAAAGACGAACAATAAGACGCTGAAAAGCGTCTTTTTTCGTTTTGGTCC |
| ECK120033736[10] | AACGCATGAGAAAGCCCCCGGAAGATCACCTTCCGGGGGCTTTTTTATTGCGC |
| **Ribozymes** | |
| RiboJ[11] | AGCTGTCACCGGATGTGCTTTCCGGTCTGATGAGTCCGTGAGGACGAAACAGCCTCTACAAATAATTTTGTTTAA |
| ElvJ[11] | AGCCCCATAGGGTGGTGTGTACCACCCCTGATGAGTCCAAAAGGACGAAATGGGGCCTCTACAAATAATTTTGTTTAA |
| PlmJ[11] | AGTCATAAGTCTGGGCTAAGCCCACTGATGAGTCGCTGAAATGCGACGAAACTTATGACCTCTACAAATAATTTTGTTTAA |
| RiboJ64[11] | AGGAGTCAATTAATGTGCTTTTAATTCTGATGAGACGGTGACGTCGAAACTCCCTCTACAAATAATTTTGTTTAA |
| **Genes** | |

| | |
|---|---|
| *tetR*<br>(codon optimized for MR-1) | ATGTCTCGTTTAGATAAATCTAAAGTTATCAACTCTGCTTTAGAATTATTAAAC<br>GAAGTTGGTATCGAAGGTTTAACTACTCGTAAATTAGCTCAAAAATTAGGTGT<br>TGAACAGCCCACATTATACTGGCACGTTAAAAACAAGAGGGCGTTATTAGAT<br>GCTCTCGCTATCGAAATGTTAGATCGTCACCACACTCACTTCTGTCCATTAGA<br>AGGTGAATCTTGGCAAGATTTCTTACGTAACAACGCTAAATCGTTCCGTTGT<br>GCGTTATTATCGCACCGTGATGGTGCTAAAGTTCACTTAGGTACTCGTCCAA<br>CTGAAAAACAATACGAAACTTTAGAAAACCAATTAGCTTTCTTATGTCAACAA<br>GGTTTCTCGCTCGAAAACGCGCTCTATGCGTTATCGGCTGTTGGCCACTTCA<br>CTTTAGGTTGTGTTTTAGAAGATCAAGAACACCAAGTTGCTAAAGAAGAACGT<br>GAAACTCCAACTACTGATTCTATGCCACCATTATTACGTCAAGCTATCGAATT<br>ATTCGATCACCAAGGTGCCGAGCCAGCGTTCCTCTTCGGTTTAGAATTAATC<br>ATCTGTGGTTTAGAAAAACAATTAAAATGTGAATCTGGTTCTTAA |
| *lacI* | GTGAAACCAGTAACGTTATACGATGTCGCAGAGTATGCCGGTGTCTCTTATC<br>AGACCGTTTCCCGCGTGGTGAACCAGGCCAGCCACGTTTCTGCGAAAACGC<br>GGGAAAAAGTGGAAGCGGCGATGGCGGAGCTGAATTACATTCCCAACCGCG<br>TGGCACAACAACTGGCGGGCAAACAGTCGTTGCTGATTGGCGTTGCCACCT<br>CCAGTCTGGCCCTGCACGCGCCGTCGCAAATTGTCGCGGCGATTAAATCTC<br>GCGCCGATCAACTGGGTGCCAGCGTGGTGGTGTCGATGGTAGAACGAAGC<br>GGCGTCGAAGCCTGTAAAGCGGCGGTGCACAATCTTCTCGCGCAACGCGTC<br>AGTGGGCTGATCATTAACTATCCGCTGGATGACCAGGATGCCATTGCTGTG<br>GAAGCTGCCTGCACTAATGTTCCGGCGTTATTTCTTGATGTCTCTGACCAGA<br>CACCCATCAACAGTATTATTTTCTCCCATGAAGACGGTACGCGACTGGGCGT<br>GGAGCATCTGGTCGCATTGGGTCACCAGCAAATCGCGCTGTTAGCGGGCCC<br>ATTAAGTTCTGTCTCGGCGCGTCTGCGTCTGGCTGGCTGGCATAAATATCTC<br>ACTCGCAATCAAATTCAGCCGATAGCGGAACGGGAAGGCGACTGGAGTGCC<br>ATGTCCGGTTTTCAACAAACCATGCAAATGCTGAATGAGGGTATCGTTCCCA<br>CTGCGATGCTGGTTGCCAACGATCAGATGGCGCTGGGCGCAATGCGCGCC<br>ATTACCGAGTCCGGGCTGCGCGTTGGTGCGGATATCTCGGTAGTGGGATAC<br>GACGATACCGAAGACAGCTCATGTTATATCCCGCCGTTAACCACCATCAAAC<br>AGGATTTTCGCCTGCTGGGGCAAACCAGCGTGGACCGCTTGCTGCAACTCT<br>CTCAGGGCCAGGCGGTGAAGGGCAATCAGCTGTTGCCCGTGTCACTGGTG<br>AAAAGAAAAACCACCCTGGCGCCCAATACGCAAACCGCCTCTCCCCGCGCG<br>TTGGCCGATTCATTAATGCAGCTGGCACGACAGGTTTCCCGACTGGAAAGC<br>GGGCAGTGA |
| *sfgfp* | ATGCGTAAAGGCGAAGAGCTGTTCACTGGTGTCGTCCCTATTCTGGTGGAA<br>CTGGATGGTGATGTCAACGGTCATAAGTTTTCCGTGCGTGGCGAGGGTGAA<br>GGTGACGCAACTAATGGTAAACTGACGCTGAAGTTCATCTGTACTACTGGTA<br>AACTGCCGGTACCTTGGCCGACTCTGGTAACGACGCTGACTTATGGTGTTCA<br>GTGCTTTGCTCGTTATCCGGACCATATGAAGCAGCATGACTTCTTCAAGTCC<br>GCCATGCCGGAAGGCTATGTGCAGGAACGCACGATTTCCTTTAAGGATGAC<br>GGCACGTACAAAACGCGTGCGGAAGTGAAATTTGAAGGCGATACCCTGGTA<br>AACCGCATTGAGCTGAAAGGCATTGACTTTAAAGAAGACGGCAATATCCTGG<br>GCCATAAGCTGGAATACAATTTTAACAGCCACAATGTTTACATCACCGCCGA<br>TAAACAAAAAAATGGCATTAAAGCGAATTTTAAAATTCGCCACAACGTGGAG<br>GATGGCAGCGTGCAGCTGGCTGATCACTACCAGCAAAACACTCCAATCGGT<br>GATGGTCCTGTTCTGCTGCCAGACAATCACTATCTGAGCACGCAAAGCGTTC<br>TGTCTAAAGATCCGAACGAGAAACGCGATCATATGGTTCTGCTGGAGTTCGT<br>AACCGCAGCGGGCATCACGCATGGTATGGATGAACTGTACAAATGATGA |
| *mtrC* | ATGATGAACGCACAAAAATCAAAAATCGCACTGCTGCTCGCAGCAAGTGCCG<br>TCACAATGGCCTTAACCGGCTGTGGTGGAAGCGATGGTAATAACGGCAATG<br>ATGGTAGTGATGGTGGTGAGCCAGCAGGTAGCATCCAGACGTTAAACCTAG<br>ATATCACTAAAGTAAGCTATGAAAATGGTGCACCTATGGTCACTGTTTTCGCC<br>ACTAACGAAGCCGACATGCCAGTGATTGGTCTCGCAAATTTAGAAATCAAAA |

| | |
|---|---|
| | AAGCACTGCAATTAATACCGGAAGGGGCGACAGGCCCAGGTAATAGCGCTA<br>ACTGGCAAGGCTTAGGCTCATCAAAGAGCTATGTCGATAATAAAAACGGTAG<br>CTATACCTTTAAATTCGACGCCTTCGATAGTAATAAGGTCTTTAATGCTCAAT<br>TAACGCAACGCTTTAACGTTGTTTCTGCTGCGGGTAAATTAGCAGACGGAAC<br>GACCGTTCCCGTTGCCGAAATGGTTGAAGATTTCGACGGCCAAGGTAATGC<br>GCCGCAATATACAAAAAATATCGTTAGCCACGAAGTATGTGCTTCTTGCCAC<br>GTAGAAGGTGAAAAGATTTATCACCAAGCTACTGAAGTCGAAACTTGTATTTC<br>TTGCCACACTCAAGAGTTTGCGGATGGTCGCGGCAAACCCCATGTCGCCTT<br>TAGTCACTTAATTCACAATGTGCATAATGCCAACAAAGCTTGGGGCAAAGAC<br>AATAAAATCCCTACAGTTGCACAAAATATTGTCCAAGATAATTGCCAAGTTTG<br>TCACGTTGAATCCGACATGCTCACCGAGGCAAAAAACTGGTCACGTATTCCA<br>ACAATGGAAGTCTGTTCTAGCTGTCACGTAGACATCGATTTTGCTGCGGGTA<br>AAGGCCACTCTCAACAACTCGATAACTCCAACTGTATCGCCTGCCATAACAG<br>CGACTGGACTGCTGAGTTACACACAGCCAAAACCACCGCAACTAAGAACTTG<br>ATTAATCAATACGGTATCGAGACTACCTCGACAATTAATACCGAAACTAAAGC<br>AGCCACAATTAGTGTTCAAGTTGTAGATGCGAACGGTACTGCTGTTGATCTC<br>AAGACCATCCTGCCTAAAGTGCAACGCTTAGAGATCATCACCAACGTTGGTC<br>CTAATAATGCAACCTTAGGTTATAGTGGCAAAGATTCAATATTTGCAATCAAA<br>AATGGAGCTCTTGATCCAAAAGCTACTATCAATGATGCTGGCAAACTGGTTT<br>ATACCACTACTAAAGACCTCAAACTTGGCCAAAACGGCGCAGACAGCGACA<br>CAGCATTTAGCTTTGTAGGTTGGTCAATGTGTTCTAGCGAAGGTAAGTTTGTA<br>GACTGTGCAGACCCTGCATTTGATGGTGTTGATGTAACTAAGTATACCGGCA<br>TGAAAGCGGATTTAGCCTTTGCTACTTTGTCAGGTAAAGCACCAAGTACTCG<br>CCACGTTGATTCTGTTAACATGACAGCCTGTGCCAATTGCCACACTGCTGAG<br>TTCGAAATTCACAAAGGCAAACAACATGCAGGCTTTGTGATGACAGAGCAAC<br>TATCACACACCCAAGATGCTAACGGTAAAGCGATTGTAGGCCTTGACGCATG<br>TGTGACTTGTCATACTCCTGATGGCACCTATAGCTTTGCCAACCGTGGTGCG<br>CTAGAGCTAAAACTACACAAAAAACACGTTGAAGATGCCTACGGCCTCATTG<br>GTGGCAATTGTGCCTCTTGTCACTCAGACTTCAACCTTGAGTCTTTCAAGAA<br>GAAAGGCGCATTGAATACTGCCGCTGCAGCAGATAAAACAGGTCTATATTCT<br>ACGCCGATCACTGCAACTTGTACTACCTGTCACACAGTTGGCAGCCAGTACA<br>TGGTCCATACGAAAGAAACCCTGGAGTCTTTCGGTGCAGTTGTTGATGGCAC<br>AAAAGATGATGCTACCAGTGCGGCACAGTCAGAAACCTGTTTCTACTGCCAT<br>ACCCCAACAGTTGCAGATCACACTAAAGTGAAAATGTAA |
| *mtrA* | ATGAAGAACTGCCTAAAAATGAAAAACCTACTGCCGGCACTTACCATCACAA<br>TGGCAATGTCTGCAGTTATGGCATTAGTCGTCACACCAAACGCTTATGCGTC<br>GAAGTGGGATGAGAAAATGACGCCAGAGCAAGTCGAAGCCACCTTAGATAA<br>GAAGTTTGCCGAAGGCAACTACTCCCCTAAAGGCGCCGATTCTTGCTTGATG<br>TGCCATAAGAAATCCGAAAAAGTCATGGACCTTTTCAAAGGTGTCCACGGTG<br>CGATTGACTCCTCTAAGAGTCCAATGGCTGGCCTGCAATGTGAGGCATGCC<br>ACGGCCCACTGGGTCAGCACAACAAAGGCGGCAACGAGCCGATGATCACTT<br>TTGGTAAGCAATCAACCTTAAGTGCCGACAAGCAAAACAGCGTATGTATGAG<br>CTGTCACCAAGACGATAAGCGTATGTCTTGGAATGGCGGTCACCATGACAAT<br>GCCGATGTTGCTTGTGCTTCTTGTCACCAAGTACACGTCGCAAAAGATCCTG<br>TGTTATCTAAAAACACGGAAATGGAAGTCTGTACTAGCTGCCATACAAAGCA<br>AAAAGCGGATATGAATAAACGCTCAAGTCACCCACTCAAATGGGCACAAATG<br>ACCTGTAGCGACTGTCACAATCCCCATGGGAGCATGACAGATTCCGATCTTA<br>ACAAGCCTAGCGTGAATGATACCTGTTATTCCTGTCACGCCGAAAAACGCGG<br>CCCAAAACTTTGGGAGCATGCACCCGTCACTGAGAATTGTGTCACTTGCCAC<br>AATCCTCACGGTAGTGTGAATGACGGTATGCTGAAAACCCGTGCGCCACAG<br>CTATGTCAGCAATGTCACGCCAGCGATGGCCACGCCAGCAACGCCTACTTA<br>GGTAACACTGGATTAGGTTCAAATGTCGGTGACAATGCCTTTACTGGTGGAA |

| | |
|---|---|
| | GAAGCTGCTTAAATTGCCATAGTCAGGTTCATGGTTCTAACCATCCATCTGG CAAGCTATTACAGCGCTAA |
| *cymA* | ATGAACTGGCGTGCACTATTTAAACCCAGCGCGAAATATTCCATCCTAGCGC TACTGGTTGTTGGTATCGTGATTGGTGTTGTGGGCTATTTTGCAACTCAGCA GACTTTACATGCGACAAGTACAGATGCGTTCTGTATGTCTTGCCATAGCAAT CATTCCTTGAAGAATGAAGTGCTGGCATCTGCCCACGGTGGCGGCAAAGCC GGGGTTACTGTTCAGTGTCAAGACTGTCACTTACCCCATGGCCCTGTTGATT ATTTAATTAAGAAAATCATCGTATCTAAAGATTTATATGGTTTCTTAACTATTG ATGGCTTTAACACTCAAGCTTGGTTAGACGAAAACCGCAAAGAGCAAGCCGA CAAAGCATTGGCTTACTTCCGTGGTAACGACTCAGCAAACTGTCAACACTGC CATACTCGCATTTATGAAAACCAGCCAGAAACCATGAAGCCAATGGCTGTGA GAATGCACACCAACAACTTCAAGAAAGATCCTGAAACGAGAAAGACCTGTGT GGATTGCCACAAAGGTGTCGCTCACCCCTATCCAAAAGGATAA |

**Table S5.** Nonlinear fit parameters for main text Figures 4-6

| **Figure** | **Fitted Values** | | | | **95% Confidence Interval** | | | | **$R^2$ (weighted)** |
|---|---|---|---|---|---|---|---|---|---|
| | MIN | n | MAX | $K_{1/2}$ | MIN | n | MAX | $K_{1/2}$ | |
| Figure 4b | 6124 | 1.699 | 29301 | 97.83 | 5698 to 6715 | 1.317 to 2.118 | 27860 to 32435 | 82.29 to 124.2 | 0.9559 |
| Figure 4c | 0.6299 | 1.296 | 2.578 | 14.2 | 0.5184 to 0.8713 | 0.7638 to 2.463 | 2.445 to 3.150 | 9.113 to 30.58 | 0.8232 |
| Figure 4d | = 0.2767 | 0.653 | = 1.318 | 221.4 | n.d. | 0.4771 to 1.365 | n.d. | 37.05 to 282.2 | 0.5958 |
| Figure 4e | 0.2731 | 1.75 | 1.368 | 15.53 | 0.2087 to 0.4044 | 0.8473 to 3.638 | 1.295 to 1.731 | 10.29 to 28.25 | 0.8385 |
| Figure 5b | 15.83 | 1.176 | 113.2 | 79.54 | 10.38 to 20.84 | 0.9112 to 1.521 | 107.6 to 121.0 | 64.82 to 98.42 | 0.9676 |
| Figure 5c | 0.8899 | 1.199 | 37.6 | 279.4 | 0.8290 to 1.394 | 1.046 to 1.519 | 31.24 to 46.20 | 187.7 to 428.5 | 0.9499 |
| Figure 5d | 0.7018 | 0.6978 | 35.63 | 188.1 | 0.4938 to 1.515 | 0.4875 to 0.9073 | 30.18 to 235.1 | 90.48 to 25541 | 0.8319 |
| Figure 6b | = 0.6370 | 2.419 | = 3.075 | 50.44 | n.d. | 1.246 to n.d. | n.d. | 45.34 to 248.2 | 0.7014 |
| Figure 6c | 0.4392 | 1.93 | 2.27 | 17.46 | 0.3645 to 1.439 | 0.3533 to 3.380 | 1.666 to 3.560 | 5.951 to 68.18 | 0.5635 |
